# PaNDA: Efficient Optimization of Phylogenetic Diversity in Networks

**DOI:** 10.1101/2025.11.14.688467

**Authors:** Niels Holtgrefe, Leo van Iersel, Ruben Meuwese, Yukihiro Murakami, Jannik Schestag

## Abstract

Phylogenetic diversity plays an important role in biodiversity, conservation, and evolutionary studies by measuring the diversity of a set of taxa based on their phylogenetic relationships. In phylogenetic trees, a subset of *k* taxa with maximum phylogenetic diversity can be found by a simple and efficient greedy algorithm. However, this algorithmic tractability is lost when considering phylogenetic networks, which incorporate reticulate evolutionary events such as hybridization and horizontal gene transfer. To address this challenge, we introduce `PaNDA` (Phylogenetic Network Diversity Algorithms), the first software package and interactive graphical user-interface for exploring, visualizing and maximizing diversity in phylogenetic networks. `PaNDA` includes a novel algorithm to find a subset of *k* taxa with maximum diversity, running in polynomial time for networks of bounded *scanwidth*, a measure of tree-likeness of a network that grows slower than the well-known *level* measure. This algorithm considers the variant of phylogenetic diversity on networks in which the branch lengths of all paths from the root to the selected taxa contribute towards their diversity. We demonstrate the scalability of this algorithm on simulated networks, successfully analyzing level-15 networks with up to 200 taxa in seconds. We also provide a proof-of-concept analysis using a phylogenetic network on *Xiphophorus* species, illustrating how the tool can support diversity studies based on real genomic data. The software is easily installable and freely available at https://github.com/nholtgrefe/panda. Additionally, we extend the definition of phylogenetic diversity to semi-directed phylogenetic networks, which are mixed graphs increasingly used in phylogenetic analysis to model uncertainty of the root location. We prove that finding a subset of *k* taxa with maximum diversity remains NP-hard on semi-directed networks, but do present a polynomial-time algorithm for networks with bounded level.

## 1 Introduction

A prominent measure of biodiversity of a set of species, or taxa, is their *phylogenetic diversity* [13], which has found widespread application in conservation [16, 17, 23], biogeography [36], and human health [4, 31]. Defined as the total branch length of the subtree induced by the considered taxa in a phylogenetic tree, phylogenetic diversity serves as a key indicator of heterogeneity and evolutionary remoteness [8]. In part, phylogenetic diversity has earned its popularity because the associated optimization problem, in which a subset of *k* taxa with maximum phylogenetic diversity is determined, can be solved efficiently by a greedy algorithm [33, 41].

Phylogenetic networks, which generalize phylogenetic trees by accounting for reticulate evolutionary events such as hybridization and horizontal gene transfer, offer a more realistic view of the evolutionary history of taxa in many cases [3, 29]. With the rise of methods that infer phylogenetic networks [29], the natural question has been raised: How should phylogenetic diversity be measured in phylogenetic networks? One intuitive measure is *all-paths phylogenetic diversity* [46], which generalizes the definition on trees by considering the total length of branches on all paths from the root to the considered set of taxa. However, accommodating reticulate events comes at a computational cost: the associated maximization problem Maximize-All-Paths-PD (MapPD) (see Section 2.1 for definitions) is NP-hard [7].

Although some algorithms have been presented for MapPD [7, 24, 25], these are of a theoretical nature and have not been implemented. Also, other variants of phylogenetic diversity on networks have recently received much interest from a theoretical point of view [7, 24, 25, 42, 43, 44]. For example, it has been shown that for the more advanced average-tree phylogenetic diversity model, in which the diversity score is taken as the weighted average over all displayed trees, even computing the diversity score for a fixed set of taxa is computationally hard (more precisely, it is #P-hard) [42]. Instead, MapPD strikes a better balance between biological accuracy and reasonable computationally complexity. Empirical work on phylogenetic diversity in networks has been restricted to [46], but its brute-force approach makes it only applicable to very small data sets. Related practical work was also presented in [9, 45], but these consider different types of input: trees with feature presence-absence data [9] or unrooted data-display networks [45]. None of these includes an easy-to-use and scalable software tool.

To address this gap, we introduce `PaNDA` (Phylogenetic Network Diversity Algorithms): an interactive software package for diversity optimization, visualization, and exploration in phylogenetic networks. `PaNDA` contains a novel algorithm that optimally solves MapPD in polynomial time on networks of bounded *scanwidth* [6], a measure of tree-likeness that is particularly suitable for designing practical algorithms for phylogenetic networks. For example, networks with small scanwidth may still have large *level* [22], a well-known measure of network tree-likeness defined as the maximum number of edges that needs to be deleted per *blob* (reticulated or biconnected component) to turn the network into a tree. The algorithm has running time 𝒪 (2^sw^ sw *k*^2^*m*), with sw the scanwidth, *m* the number of edges and *k* the number of taxa to select. This excludes the time necessary to compute the scanwidth, which can be done efficiently in practice [22] and was fast enough for all considered instances.

To make diversity analysis in networks accessible and user-friendly, `PaNDA` is supplied with a graphical user-interface (GUI) that facilitates both optimization and exploration, see Figure 1. Through this interface, users can use the underlying MapPD-algorithm to find subsets of taxa of maximum phylogenetic diversity, interactively explore how diversity changes across different combinations of taxa, and visualize networks. Moreover, `PaNDA` provides an excellent framework to which other variants can be added in future work, which will make it easier to compare and combine these variants in benchmark studies and biological applications.

**Figure 1.**
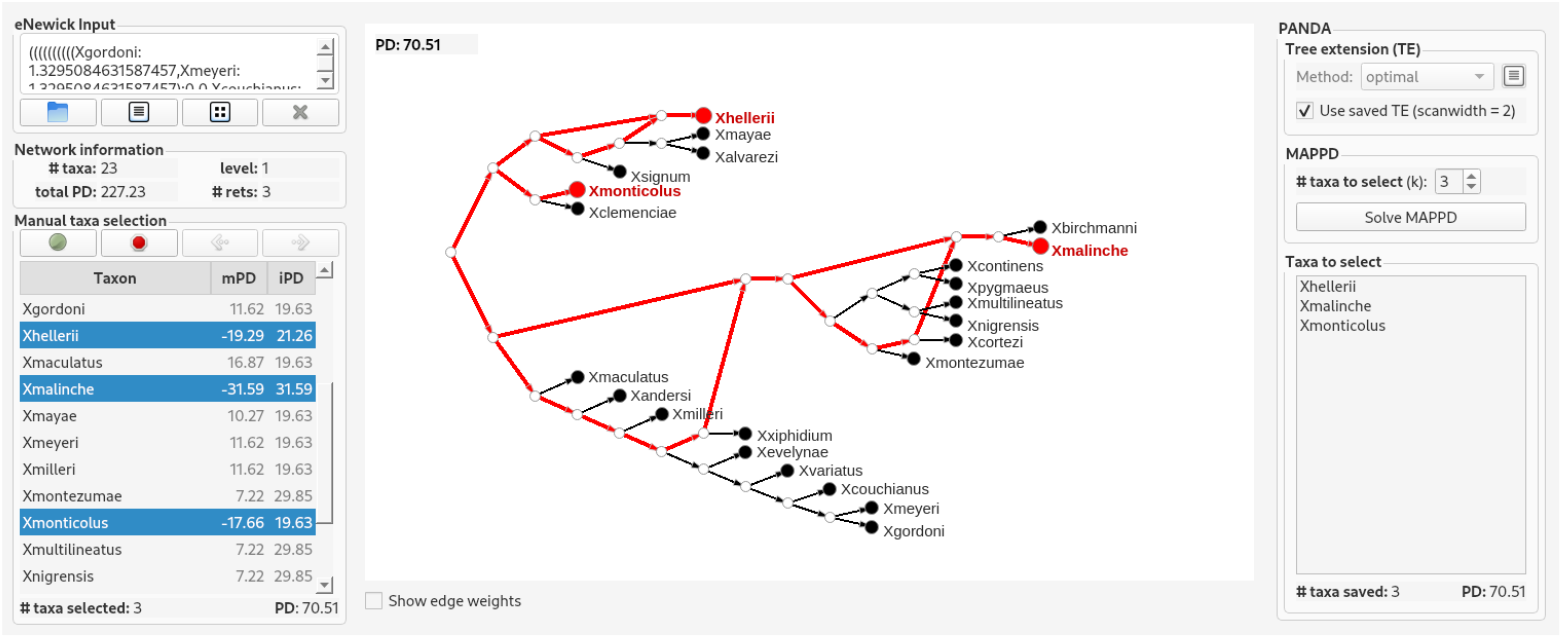
The `PaNDA` graphical user-interface with the network presented in Figure 6 as input.

We evaluate the performance of our algorithm through extensive simulations and show that it scales well, even on networks far beyond the size and complexity currently supported by existing network inference methods [29]. Specifically, optimal solutions on level-15 phylogenetic networks with up to 200 leaves can be found in seconds. To demonstrate its real-world applicability, we use `PaNDA` to study a phylogenetic network on a set of *Xiphophorus* species, showcasing how it can be used to meaningfully analyze empirical data and support biologically relevant insights.

In addition to this practical contribution, we introduce a new variant of the MapPD problem where the input network is a *semi-directed* phylogenetic network which contains both directed and undirected edges. These networks have gained attention recently as they account for uncertainty in root placement, addressing unidentifiability issues that arise under several evolutionary models [2, 15]. We show that MapPD on semi-directed networks remains NP-hard and we present an algorithm solving it in 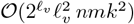 time, with *n* the total number of taxa and *ℓ*_*v*_ the *visible vertex level*, a parameter that can be much smaller than the level. The main advantage of this algorithm is that it can be applied directly to semi-directed networks produced by statistically consistent inference methods [1, 19, 39]. Both presented algorithms are applicable to binary as well as to nonbinary networks.

## 2 Methods

### 2.1 Preliminaries

For *k* ∈ ℕ_*>*0_, we write [*k*] = {1, …, *k*} and [*k*]_0_ = {0, 1, …, *k*}. We use standard graph notations [12].

#### Graphs

Throughout this paper, we consider only simple mixed graphs, i.e., graphs that may contain both directed and undirected edges but no multiple edges between the same pair of vertices. Given a mixed graph *G*, for *U* ⊆ *V* (*G*), we denote by *G*[*U*] the *subgraph induced by U*. Given a vertex *v* ∈ *V* (*G*), we use 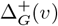 to denote its set of outgoing directed edges and 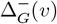 for its set of incoming directed edges, where we omit the subscript if *G* is clear from context.

We consider four types of paths in a mixed graph, all of which are assumed to be vertex-disjoint. A path is an *edge path* if it only contains undirected edges. A path is *semi-directed* if all its directed edges are oriented in the same direction. A *directed path* is a semi-directed path that does not contain undirected edges. An *up-down path P* = *v*_0_ … *v*_*t*_ between two vertices *v*_0_ and *v*_*t*_, which are called *endpoints*, satisfies that both *v*_*i*_ … *v*_0_ and *v*_*i*_ … *v*_*t*_, for some *i* ∈ [*t*], are semi-directed paths. We note that every edge path is a semi-directed path, every directed path is a semi-directed path, every semi-directed path is an up-down path, but the converse do not hold.

#### Phylogenetic networks

A *directed (phylogenetic) network* 𝒩^+^ on a set of at least two taxa *X* is a directed acyclic graph such that (i) there is a unique vertex with in-degree zero (the *root*) and its out-degree is at least two; (ii) the vertices with out-degree zero (the *leaves*) have in-degree one and they are bijectively labeled by elements of *X*; (iii) all other vertices either have in-degree one and out-degree at least two (*tree vertices*), or in-degree at least two and out-degree one (*reticulation vertices*, or simply *reticulations*). See Figure 2a(i) for an example. The edges entering a reticulation vertex are called *reticulation edges*.

**Figure 2.**
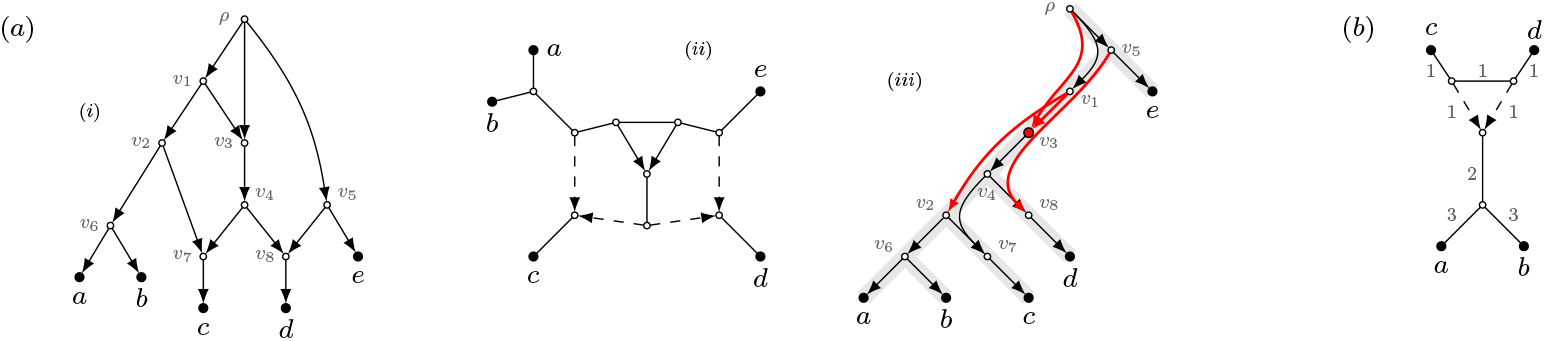
(a): A directed phylogenetic network on *X* = {*a, b, c, d, e* } (i), its associated semi-directed network (ii), and a tree-extension of the directed network with a scanwidth of 4 (iii). The edges that make up the tree extension are drawn in gray, with the edges of the original graph drawn on top of it in black and red. The scanwidth of 4 is attained at the vertex *v*_3_ (highlighted), since the set GW(*v*_3_) contains 4 edges (in thick red). (b): A semi-directed network on {*a, b, c, d*} with edge-weights illustrating that naively iterating over all rootings of a semi-directed network and solving MapPD does not solve MapSPD (see text).

A *semi-directed (phylogenetic) network* 𝒩^−^ on a set of at least two taxa *X* is a mixed graph which can be obtained from a directed network 𝒩^+^ on *X* by replacing every edge with an undirected edge except for the reticulation edges, and subsequently suppressing the former root vertex if its degree is 2. See Figure 2a(ii). Note that we may still refer to the leaves, tree vertices, reticulation vertices and reticulation edges of a semi-directed network. The reticulation edges are precisely the directed edges. We say that a vertex *u* is *above* a non-reticulation vertex *v* if there exists a semi-directed path from *u* to *v*. We say that a vertex *u* is *above* a reticulation vertex *v* if there exists a semi-directed path that is not an edge path from *u* to *v*.

Throughout this paper, we assume that every (directed or semi-directed) network 𝒩 is equipped with an *edge weight* function *ω*: *E*(𝒩) → ℝ_≥0_, which may, e.g., represent branch lengths. We generalize *ω* to subsets *B* ⊆ *E*(𝒩) by *ω*_Σ_(*B*) = ∑_*e*∈*B*_ *ω*(*e*). We call 𝒩 *binary* if the degree of each vertex is either one or three, with the exception of a possible root which has degree two. Note that the directed network in Figure 2a is not binary, but that the semi-directed network in Figure 2b is binary. A *blob B* of 𝒩 is a maximal subgraph without any cut-edges, and it is *non-trivial* if it contains at least one reticulation vertex. As an example, the vertices {*ρ, v*_1_, *v*_2_, *v*_3_, *v*_4_, *v*_5_, *v*_7_, *v*_8}_ form the vertex set of a non-trivial blob of the network in Figure 2a.

A reticulation *r* is *visible* if there is a leaf *x* such that for every vertex *u* above *r*, every semi-directed path from *u* to *x* passes through *r*. The *visible vertex level* of is the maximum number of 𝒩 visible reticulations in any of its blobs, and the *vertex level* of is the maximum number of reticulations in any of 𝒩 its blobs. For binary networks, the vertex level equals the standard notion of the *level* of a network: the maximum number of reticulation edges to be deleted in any of its blobs to turn the network into a tree. For example, in the network of Figure 2a, the reticulations *v*_7_ and *v*_8_ are visible, but the reticulation *v*_3_ is not. Hence, the visible vertex level of that network is 2, whereas both its level and vertex level are 3.

We reserve the letters *n* and *m* for the number of leaves and edges in a network, respectively.

#### Phylogenetic diversity

A *root-leaf path* in a directed network 𝒩^+^ on *X* is a directed path from the root of 𝒩^+^ to a leaf in *X*. A leaf *x* ∈ *X* is in the *offspring* off(*e*) of an edge *e*, if there is a root-leaf path to *x* that traverses *e*. Given a set of leaves *Y* ⊆ *X*, we use the notation RP(*Y*) to denote the set of all edges on root-leaf paths towards leaves in *Y*. The *all-paths phylogenetic diversity* 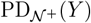 *of Y* is then

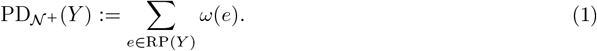

We now formally define MapPD, introduced and proved to be NP-hard even for binary networks in [7].

Maximize-All-Paths-PD (Mappd)

**Input:** A directed network 𝒩^+^ on *X* with edge-weight function *ω*; integers *k, D*.

**Question:** Is there a set *Y* ⊆ *X* of size at most *k*, such that 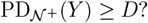

Since semi-directed networks have no roots by definition, we define a new phylogenetic diversity score for a semi-directed network 𝒩^−^ on *X* based on up-down paths. Given a set of vertices *Y*, we use the notation UP(*Y*) to denote the set of all edges on up-down paths between any pair of vertices in *Y*. Let *Y* ⊆ *X* be a set of leaves. The *all-paths semi-directed phylogenetic diversity* 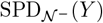 *of Y* is then

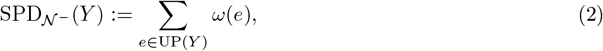

where we say that an edge *uv contributes* to 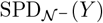 if *uv* ∈ UP(*Y*). For example, letting 𝒩^−^ be the network in Figure 2(b), 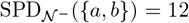 and 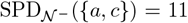. Observe that SPD has the attractive property that it is a generalization of the standard definition of phylogenetic diversity for unrooted phylogenetic trees [13]. With this new notion, we can define the new problem MapSPD.

Maximize-All-Paths-SPD (Mapspd)

**Input:** A semi-directed network 𝒩 ^−^ on *X* with edge-weight function *ω*; integers *k, D*.

**Question:** Is there a set *Y* ⊆ *X* of size at most *k*, such that 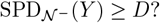

Mappd can be reduced to Mapspd by adding a high-weight edge from the root to a new leaf and then making the network semi-directed. This results in Theorem 1.

##### Theorem 1

Mapspd *is* NP*-hard, even for binary semi-directed networks*.

Note that the reverse direction is not as obvious as it may seem: naively iterating over all possible roots of a semi-directed network and solving MapPD does not solve MapSPD. Consider, for example, the network in Figure 2(b) with *k* = 2. In any rooting of the network the root is forced to be above the reticulation vertex, always having {*a, b*} as optimal MapPD solution with a value of 11. Contrary, any optimal solution for MapSPD consists of one leaf from {*a, b*} and one from {*c, d*}, yielding an optimal value of 9. So, no rooting of the network has an optimal MapPD solution corresponding to an optimal MapSPD solution in the semi-directed network.

We note that we state the problems as decision problems, but in our empirical study we in fact solve the optimization variants. That is, we find a set of taxa that maximizes the all-path phylogenetic diversity over all sets of size *k*.

#### Scanwidth

Given a directed acyclic graph *G*, we define a relation ≻_*G*_ on *V* (*G*) where *u* ≻_*G*_ *v* if there exists a directed path from *u* to *v* in *G*. We extend this relation to ⪰_*G*_, where *u* ⪰_*G*_ *v* if either *u* ≻_*G*_ *v* or *u* = *v*. A *tree extension* of a directed acylic graph *G* is a rooted tree 𝒯_*G*_ with *V* (𝒯_*G*_) = *V* (*G*) and such that *u* ≻_*G*_ *v* implies 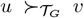 for all *u, v* ∈ *V* (*G*). For every vertex *v* ∈ *V* (𝒯_*G*_), we let 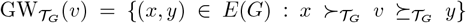, where we often omit subscripts if *G* and 𝒯_*G*_ are clear from context. Note that the sets GW(*v*) contain edges of the original graph, but the succession relations are according to the given tree extension. The *scanwidth* 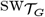 of the tree extension 𝒯_*G*_ is 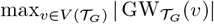 and we call 𝒯_*G*_ *optimal* if it has the smallest scanwidth among all tree extensions of *G*. The *scanwidth* sw_*G*_ of a directed acyclic graph *G*, first introduced in [6], is the scanwidth of an optimal tree extension of *G*. See Figure 2a(iii) for an example. The subtree of 𝒯_𝒩_ rooted at *v* is denoted 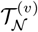.

### 2.2 Algorithm for directed networks

We present Algorithm 1, which solves an instance (𝒩, *X, ω, k, D*) of MapPD in polynomial time for directed networks of bounded scanwidth. The algorithm considers a tree-extension 𝒯_𝒩_ in a bottom-up fashion and uses a dynamic programming table to store intermediate solutions. We utilize a *binary resolution* of the network to handle cases where the input network is nonbinary. This is outlined in Lemma 2. The time complexity and correctness of the algorithm are proved in Theorem 2.

**Theorem 2**. *Algorithm 1 solves instances* (𝒩, *X, ω, k, D*) *of* MapPD *in* 𝒪 (2^sw^ sw ·*k*^2^*m*) *time, if a tree-extension* 𝒯_𝒩_ *of* 𝒩 *of scanwidth* sw *is given*.

The intuition of the table DP at the core of Algorithm 1 is as follows. For a vertex *v* ∈ *V* (𝒩), a set of edges Φ ⊆ GW(*v*), and an integer *t* ∈ [*k*]_0_, the table entry DP[*v*, Φ, *t*] stores *ω*_Σ_(Φ) plus the phylogenetic diversity maximized by a set *A* ⊆ *X* of size *t* in 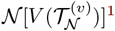^1^ where off(*e*) ∩ *A*≠ ∅ for each *e* ∈ Φ. If such a set does not exist, it stores −∞, which, in practice, can be replaced by a large negative value. When *v* is the root *ρ* of the tree-extension, Φ is empty, and *t* equals *k*, the table therefore stores the maximum phylogenetic diversity in 𝒩 of a set of *k* leaves, thereby solving MapPD. Lines 6, 8 and 10 of the algorithm cover the three base cases of the table, whereas Lines 13 and 15 contain recurrence relations that compute table entries based on entries corresponding to vertices further down in the tree.

We note that 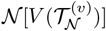 may not necessarily be a directed network, and hence we naturally extend the definition of PD_𝒩_ to general directed acyclic graphs.

### 2.3 Algorithm for semi-directed networks

In this section, we study a generalization of the all-paths semi-directed phylogenetic diversity SPD_𝒩_. Let an integer *k*, a semi-directed network 𝒩 and functions *σ*: *X* → [*k*]_0_ and *f*: *X* × [*k*]_0_ → ℝ_≥0_∪ {−∞} be given. Denote by *X*_*σ*_ the set of leaves *x* with *σ*(*x*) *>* 0. We define the *generalized phylogenetic diversity* GPD_𝒩_ (*σ*) as

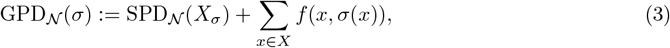

where we say that *a budget of h is spent on a set of leaves L* if ∑_*x*∈*L*_ *σ*(*x*) ≤ *h* and *σ*(*x*) *>* 0 for each *x ∈ L*. In a sense, *σ* models how much of the ‘budget’ *k* is ‘invested’ in each taxon, and *f* models additional diversity to be acquired, dependent on the choice of *σ*. We study the following problem, where one can replace −∞ with a large negative value in practice.

#### Algorithm 1

Solving Mappd parameterized by the scanwidth

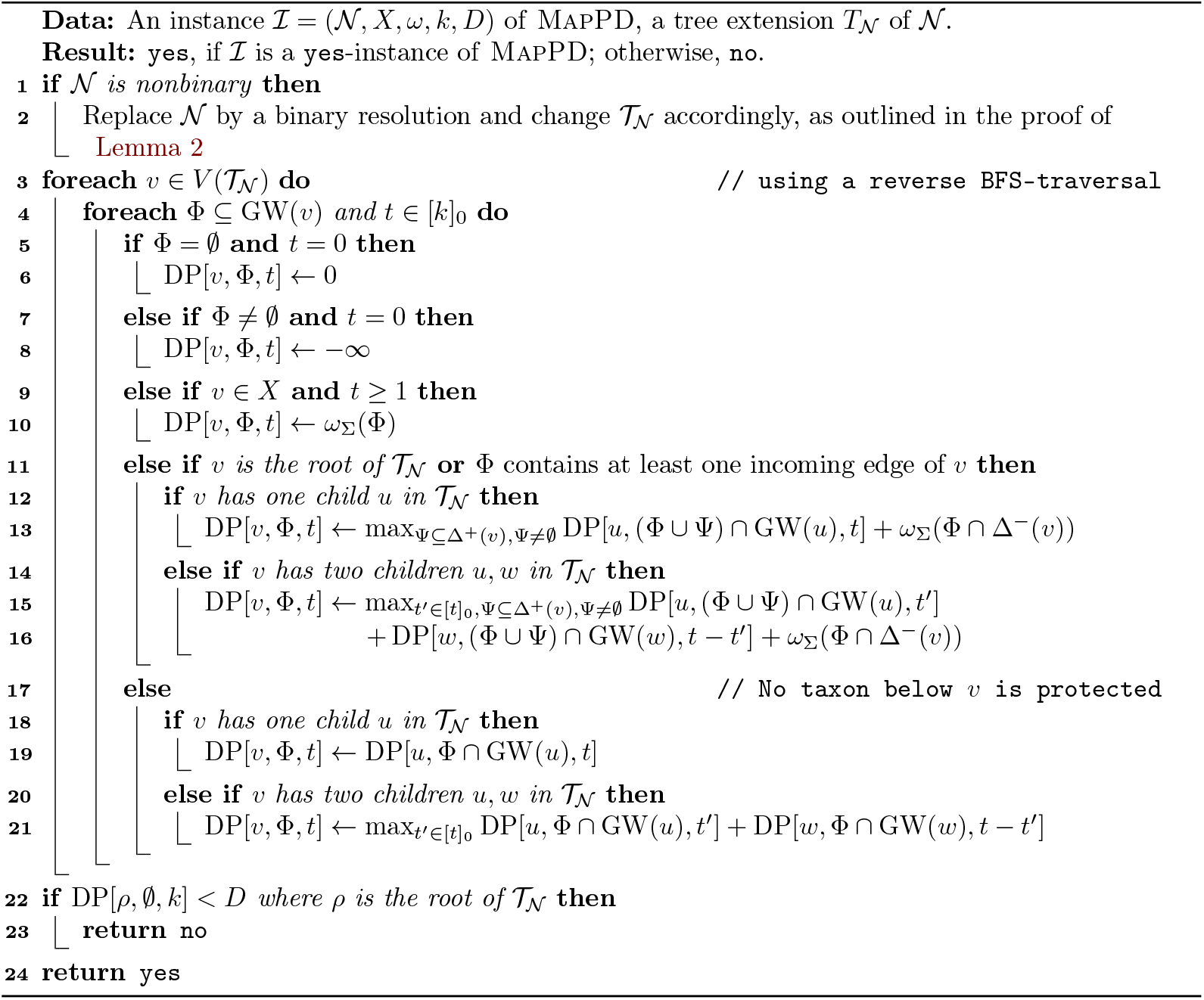

Generalized-Maximize-All-Paths-SPD (G-MapSPD)

**Input:** A semi-directed network 𝒩 on *X*, integers *k, D* ∈ ℕ_*>*0_, and functions *f*: *X* × [*k*]_0_ → ℝ_≥0_ ∪ {−∞} and *ω*: *E*(𝒩) → ℝ_≥0_.

**Question:** Is there a function *σ*: *X* → [*k*]_0_ such that ∑_*x*∈*X*_ *σ*(*x*) ≤ *k* and GPD_𝒩_ (*σ*) ≥ *D*?

Observe that Mapspdm is a special case of G-MapSPD, where *f* (*x, t*) = 0 for each *x* ∈ *X*, and *t* ∈ [*k*]_0_. Thus, G-MapSPD is NP-hard by Theorem 1. Budgeted Max-PD [34] is also a special case of G-MapSPD.^2^

We present Algorithm 2, which solves instances (𝒩, *X, f, ω, k, D*) of G-MapSPD recursively in polynomial time for networks with bounded visible vertex level. Below, we explain the steps and notation in the algorithm. The correctness and running time of the algorithm are proven in Theorem 3, with Corollary 1 following by using the reduction in the proof of Theorem 1.

#### Theorem 3

*Algorithm 2 solves instances* (𝒩, *X, f, ω, k, D*) *of* G-MapSPD *in* 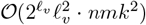 *time, where ℓ*_*v*_ *is the visible vertex level, n is the number of vertices, and m is the number of edges in* 𝒩.

#### Corollary 1

*An instance* (𝒩, *X, ω, k, D*) *of* MapPD *can be solved in* 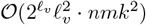 *time, where ℓ*_*v*_ *is the visible vertex level of* 𝒩.

The algorithm first uses Rule 0 to reduce all *2-blobs*: blobs with exactly 2 incident cut-edges.

#### Algorithm 2

Solving G-MapSPD parameterized by the visible vertex level

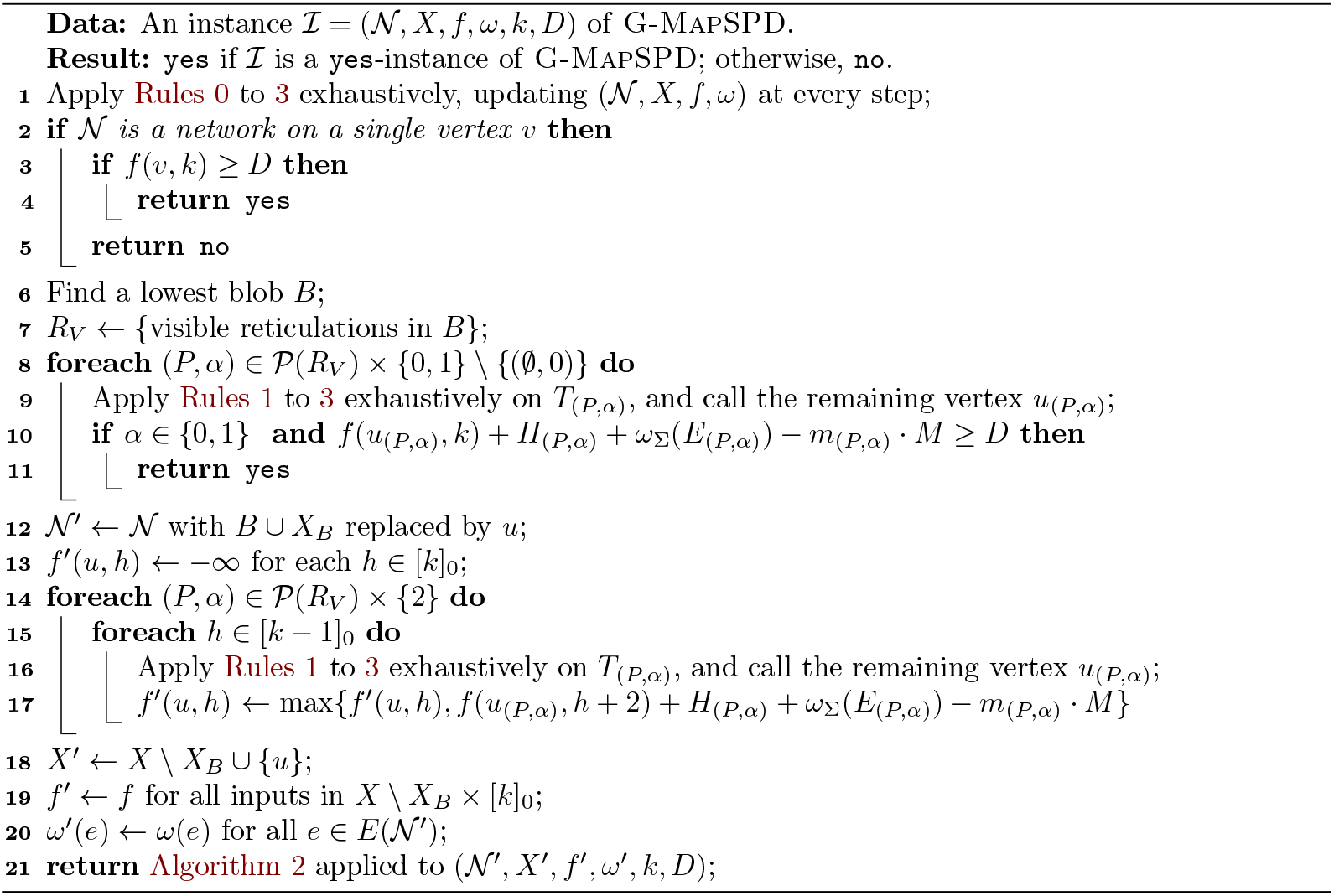

**Reduction Rule 0**. *Let B be a 2-blob with incident cut-edges* {*u,v*} *and* {*w, z*} *such that v, w* ∈ *V* (*B*). *Remove B*, {*u, v*} *and* {*w, z*} *and add the edge* {*u, z*} *with weight* ∑_*e*∈*E*(*B*)_ *ω*(*e*) + *ω*({*u, v*}) + *ω*({*w, z*}).

Algorithm 2 relies on three further reduction rules that reduce trees to single vertices. Rule 1 reduces *cherries*—two leaves connected to the same vertex—to a single leaf. Rule 2 reduces degree-two vertices that may arise from applying Rule 1. Exhaustively applying these two rules to a tree creates a single edge, which can be reduced to a single vertex with Rule 3. For integers *i, j, k*, we use *δ*_*i<j<k*_ to denote the *Kronecker delta*, i.e., *δ*_*i<j<k*_ equals 1 if *i < j < k*, and 0 otherwise.

**Reduction Rule 1**. *For given edges* {*v, x*} *and* {*v, y*} *of leaves x, y, add a new leaf z with edge* {*v, z*} *of weight* 0 *and set f* ^*′*^(*z, i*) *to* 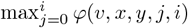 *with φ*(*v, x, y, j, i*):= *f* (*x, j*) + *f* (*y, i* − *j*) + *δ*_0*<j<k*_ · *ω*({*v, x*}) + *δ*_0*<i*−*j<k*_ · *ω*({*v, y*}) *for each i* ∈ [*k*]_0_. *Then, remove x and y with incident edges*.

Here, *f* ^*′*^ is the function for the new instance which contains *z*, but not *x* and *y*.

**Reduction Rule 2**. *Let v be a degree-2-vertex with edges* {*u, v*} *and* {*v, w*}. *Add an edge* {*u, w*} *of weight ω*({*u, v*}) + *ω*({*v, w*}) *and remove v with its incident edges*.

**Reduction Rule 3**. *Let* {*x, y*} *be the single edge of a network. Introduce a new vertex v. For each i* ∈ _0_, *set f* (*v, i*) *to* 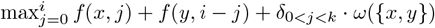. *Remove x, y and* {*x, y*}.

Before we continue, we define a non-trivial blob *B* to be a *lowest* blob, if there exists a vertex *v*_*B*_ ∈ *V* (*B*) (called a *designated vertex*), if

- all vertices that are not in *V* (*B*) and adjacent to a vertex in *V* (*B*) \{*v*_*B*_} are leaves;
- *v*_*B*_ has at most one non-leaf neighbor not in *V* (*B*); and
- *v*_*B*_ is above every vertex of *B*.

For networks containing exactly one blob, the designated vertex *v*_*B*_ can be ambiguous, but in a network of at least two blobs, every lowest blob has a unique designated vertex. The *leaf neighbors* of a blob are the leaves that are adjacent to a vertex in the blob.

If the network is not a tree, the algorithm recursively processes lowest blobs until a tree remains. Specifically, for a semi-directed network 𝒩 with vertex set *V*, the algorithm identifies a lowest blob *B* with designated vertex *v*_*B*_ on Line 6. Such a blob exists and can be found efficiently (see Lemma 8). We let *X*_*B*_ be the leaf neighbors of *B*. Algorithm 2 reduces *B* ∪ *X*_*B*_ to a single leaf *u* on Lines 12 and 18. The value *f* ^*′*^(*u, h*) for *h* ∈ [*k*]_0_—computed in the for-loop on Line 15 which is discussed below—stores the maximum generalized diversity that edges of *B* contribute when a budget of *h* is spent on leaves attached to *B*. Observe that then a budget of *k* − *h* is spent in the part above *v*_*B*_ (the graph induced by *V \*{*V* (*B*) ∪ *X*_*B*_ }).

Letting *R*_*V*_ denote the visible reticulations in *V* (*B*), the for-loop on Line 15 iterates over every set *P* in the powerset 𝒫(*R*_*V*_), and over a flag *α* ∈ {0, 1, 2}. With *P* we give a set of visible reticulations such that for each *p* ∈ *P*, some budget is forced to be spent on some leaves in 𝒞_*p*_: the set of vertices of *B* ∪ *X*_*B*_ that can be reached with an edge-path from *p*. The flag *α* indicates one of three—not necessarily distinct—options, where *α* = 0 stands for not necessarily spending budget on taxa above elements in *P, α* = 1 for definitely spending *h* ≥ 1 in 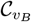, and *α* = 2 stands for definitely spending *h* ≥ 1 on some taxa of *X \X*_*B*_. For each combination of *P* and *α* (ignoring the case *P* = ∅ and *α* = 0), we then construct a tree *T*_(*P,α*)_ capturing the maximum generalized diversity that edges of *B* contribute under the scenario outlined above. On Line 9, these trees are reduced to single leaves *u*_(*P,α*)_ with Rules 1 to 3. The remaining lines in the for-loop amalgamate this information to compute the correct values of 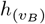.

It remains to present the construction of the trees *T*_(*P,α*)_, which requires more definitions. Fix a vertex *w* of 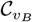 with outgoing edge *wr*, where *r* is a reticulation with a semi-directed path to a reticulation in *P*. If *P* = ∅, we set *w*:= *v*_*B*_. We define sets *P*_0_:= *P*, *P*_1_:= *P* ∪ {*w*}, and *P*_2_:= *P* ∪ {*v*_*B*_}. Let *V*_(*P,α*)_ be the vertices and *E*_(*P,α*)_ be the edges that are on up-down paths between two vertices in *P*_*α*_. In the special case that |*P* | = 1, we define *V*_(*P*,0)_ as *P* and *E*_(*P*,0)_ as ∅. In Lemma 9 we show that *V*_(*P*,1)_ and *E*_(*P*,1)_ are well-defined, regardless of the vertex *w* that is chosen. Let *M > ω*_Σ_(*E*(𝒩)) = ∑_*e*∈*E*(𝒩)_ *ω*(*e*) be a fixed integer. For a set *U* of vertices, denote by hidden(*U*) the set of vertices that can *not* be reached from *U* with an edge-path.

We define trees *T*_(*P,α*)_ by taking the graph *G*[*B* ∪ *X*_*B*_] and

1. removing hidden(*V*_(*P,α*)_) and the incident edges; let *H*_(*P,α*)_ be a the sum of *f* (*u*, 0) for each leaf in hidden(*V*_(*P,α*)_)—note that *H*_(*P,α*)_ = −∞, if *f* (*x*, 0) = −∞ for some leaf in hidden(*V*_(*P,α*)_);
2. contracting the edges *E*_(*P,α*)_ to a single vertex *v*_(*P,α*)_;
3. if *α* = 2, then adding two leaf neighbors 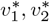 to *v* (which is identified with *v*_(*P,α*)_ if *P* ≠ ∅), setting the weight of 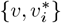 to *M*, and 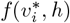 to 0 for each *h* ∈ [*k*]_0_ and each *i* ∈ {1, 2};
4. for each *r* ∈ *P*: (a) adding a vertex *v*_*r*_ and an edge {*v*_(*P,α*)_, *v*_*r*}_ of weight *M*, and (b) replacing every edge {*v*_(*P,α*)_, *w*} with *w* ∈ 𝒞_*r*_ by an edge {*v*_*r*_, *w*} of the same weight; and
5. exhaustively removing leaves not in *X*.

We define *m*_(*P,α*)_ to be the number of edges in *T*_(*P,α*)_ with a weight of *M*. In Figure 3, an example of the construction is given.

**Figure 3.**
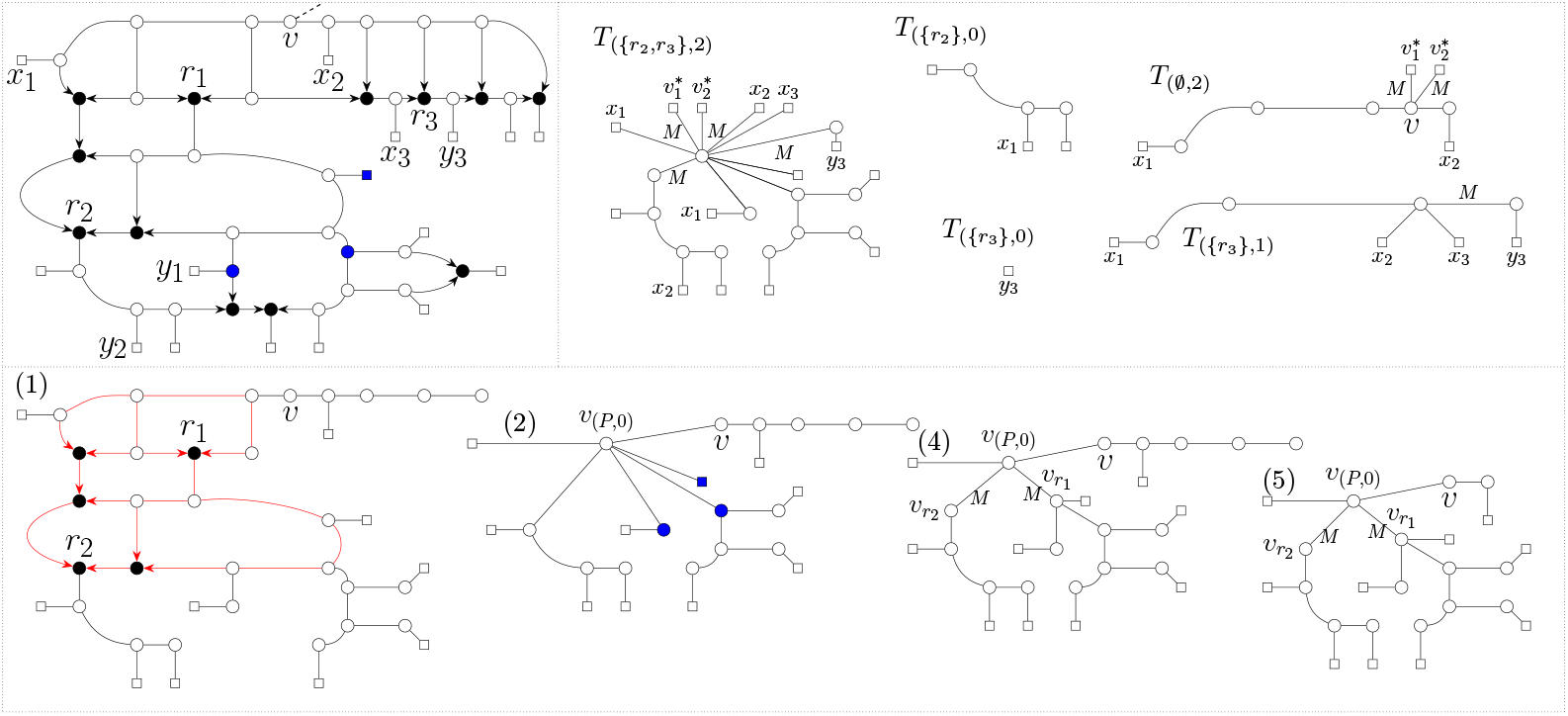
A graphic example of Algorithm 2. Observe, that reticulation *r*_*i*_ is visible, for example, with respect to *y*_*i*_, for *i* ∈ {1, 2, 3}. Not all vertices are labeled. Edge weights smaller than *M* are omitted. **Top left**: An example of a lowest blob. **Bottom**: In (*i*): The construction of *T*_(*P*,0)_ after Step *i* for *P* = {*r*_1_, *r*_2_}. In (1), the edges in *E*_(*P*,0)_—which are contracted in Step 2—are marked in red. In (2) and the top left, vertices in 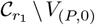 are marked in blue. **Top right**: Five other examples of constructed trees.

## 3 Software and Experiments

### Implementation

Algorithm 1 has been implemented in our `Python` package `PaNDA` in a serial manner. The algorithm takes as input a directed phylogenetic network in `eNewick` format [10] and a tree extension, which by default is computed using software originally developed in [22]. The algorithm returns—for a given integer *k*—which *k* species maximize the all-paths phylogenetic diversity and what their diversity score is. To optimize memory usage, the solutions are computed through backtracking. Additionally, we provide a graphical user interface (GUI), allowing the user to interactively explore how diversity scores vary across different sets of taxa and to visualize phylogenetic networks (see Figure 1). The package and GUI are freely available at https://github.com/nholtgrefe/panda.

### Simulations

We analyze the computational efficiency of our implementation through a simulation study, exploring how various network parameters influence its running time. Utilizing the R package `SiPhyNetwork` [26], we generated a dataset consisting of 6400 phylogenetic networks. Specifically, we exhaustively simulated networks under a birth-death-hybridization model until our data set consisted of 100 *n*-leaf level-*ℓ* networks for each combination of *n* ∈ {20, 50, 100, 200} and *ℓ* ∈ {0, …, 15}. Then, we applied our implementation of Algorithm 1 to compute the optimal phylogenetic diversity of the networks for 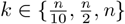.^3^ Figure 5 depicts the computation times of these experiments on a logarithmic scale. The horizontal axis represents the network level, which is polynomial-time computable and a more familiar measure of tree-likeness than scanwidth within the phylogenetic network community. Figure 4 shows how the scanwidth compares to the level in our dataset.

**Figure 4.**
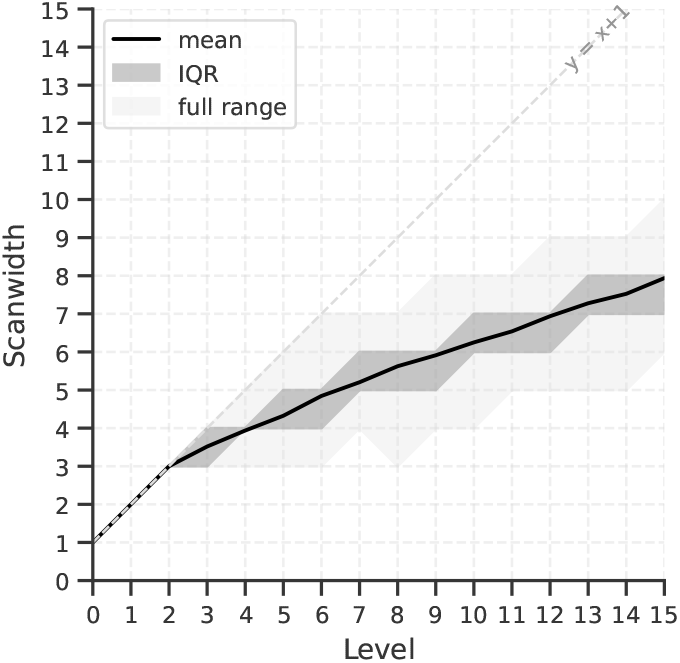
Scanwidth values plotted against the level for all networks in our dataset. The plot shows the mean, the interquartile range (IQR)—the middle 50% of the data—, and the full range of scanwidth values. The dotted line annotated ‘*y* = *x* + 1’ indicates the theoretical upper bound, showing that scanwidth is at most level+1 [22].

**Figure 5.**
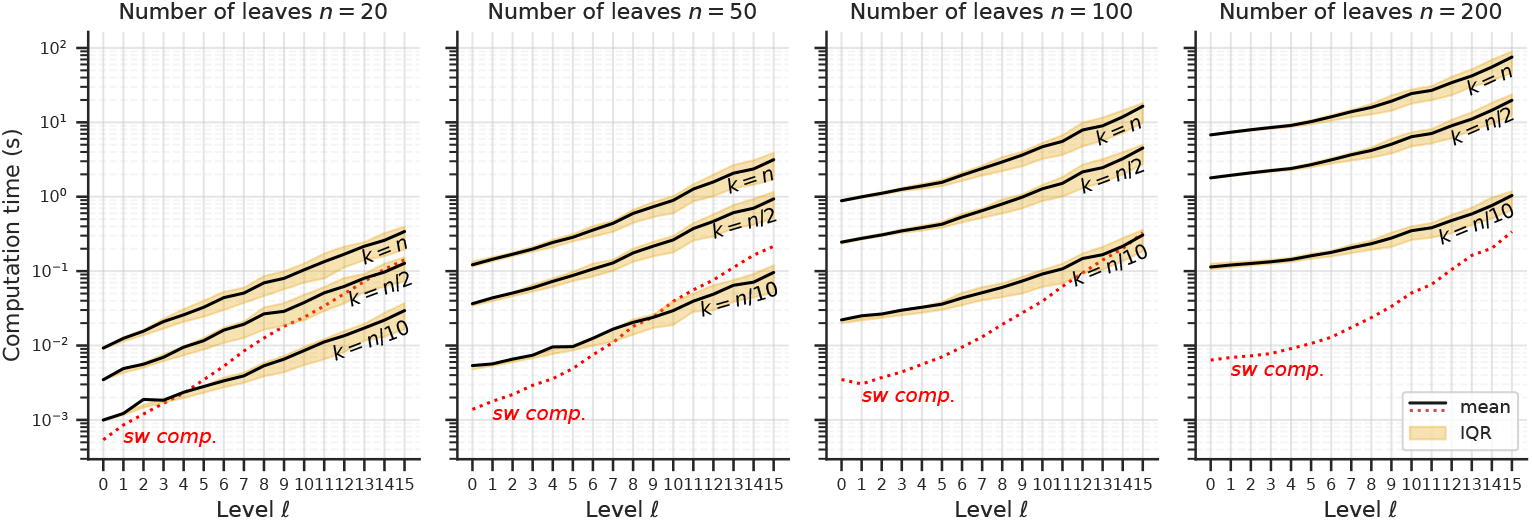
Computation times of Algorithm 1 for sets of 100 *n*-leaf level-*ℓ* networks, using *k*-values of 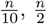 and *n*. Both the mean and interquartile range (IQR)—the middle 50% of the data—are depicted. The dotted line annotated ‘sw comp.’ shows the mean computation time for computing an optimal tree-extension using the software from [22]; this time is not included in the other measurements.

Our implementation shows great scalability, solving our largest instances in the order of seconds. Importantly, the computation of an optimal tree-extension does not pose a bottleneck for these instances, consistently completing in under one second. However, for extremely large networks, it may be advantageous to employ a heuristic tree-extension [22], which can reduce preprocessing time at the expense of increased runtime for Algorithm 1. This trade-off is particularly relevant for small values of *k*, where the tree-extension computation gets close to dominating the total runtime of Algorithm 1.

### Biological Data

To demonstrate the practical applications and capabilities of the `PaNDA` software tool, we performed a phylogenetic diversity analysis focusing on 23 *Xiphophorus* fish species consisting of three major clades: *northern swordtails, southern swordtails*, and *platyfishes*. This genus is known to have undergone extensive hybridization (see, e.g., [11]) and has thus been used as input for several phylogenetic network inference methods [5, 19, 30, 39]. We examine a level-1 phylogenetic network with branch lengths inferred in [5] with the `PhyloNetworks` package [40] (see Figure 6).

**Figure 6.**
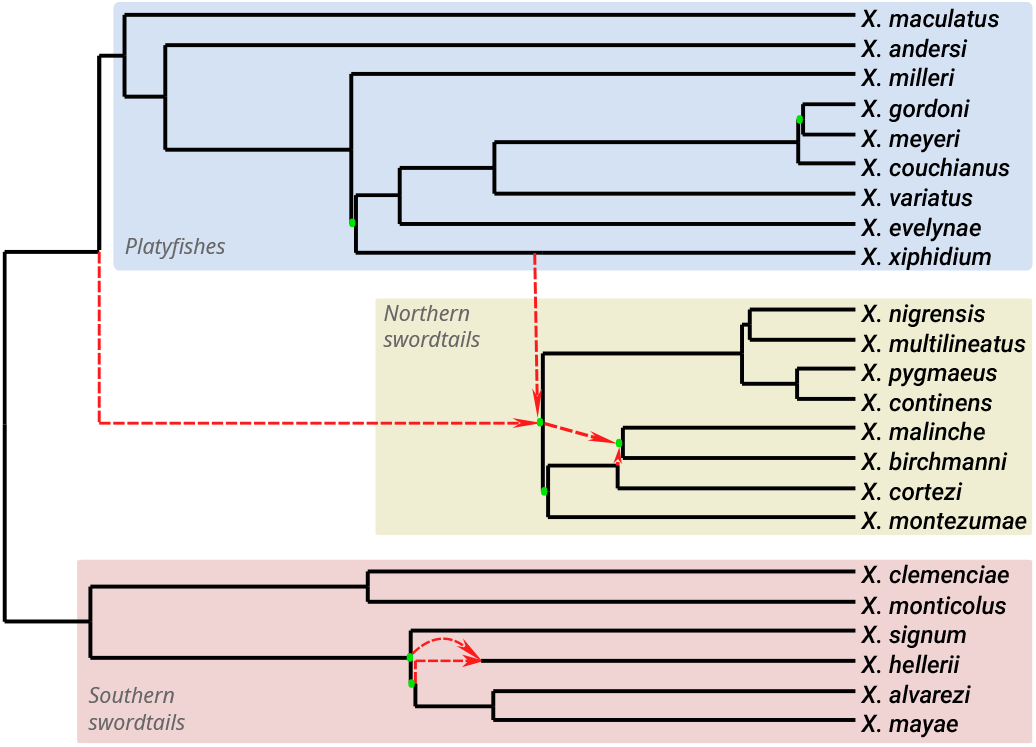
A level-1 phylogenetic network with branch lengths on 23 *Xiphophorus* fishes from [5]. The red dashed edges are reticulation edges, and branches with length zero are indicated by a green dot, to show the network topology.

Using `PaNDA`, we computed the set of *k* species that maximize the phylogenetic diversity for various values of *k*. Interestingly, for *k* = 3, the optimal solution does not include one species from each of the three traditionally defined major clades. Instead, the selected species are *X. hellerii, X. malinche*, and *X. monticolus*. While this may appear surprising at first, it is consistent with the structure of the inferred network. The inclusion of *X. hellerii* can be attributed to its hybrid origin, which allows it to capture phylogenetic signal from multiple ancestral lineages. *X. malinche* shares substantial ancestry with a large portion of the platyfishes, so no separate platyfish species is needed to represent that lineage. Furthermore, the *X. monticolus*/*X. clemenciae* lineage is essentially more phylogenetically distinct, arising from deep divergence relative to other clades. In particular, in this network, the first divergence event after the root creates the *X. monticolus*/*X. clemenciae* lineage. Together, these three species maximize phylogenetic diversity not by representing each nominal clade, but by capturing both extensive ancestral coverage and deep evolutionary distinctiveness across the network.

## 4 Discussion

In this paper, we introduced `PaNDA`, the first easy-to-use and scalable software package and graphical user interface designed to compute and optimize phylogenetic diversity (PD) in phylogenetic networks. Although currently only equipped with the all-paths phylogenetic diversity computations considered in this paper (Algorithm 1), in the future it can be extended with other algorithms, such as Algorithm 2 and algorithms for other PD-measures on phylogenetic networks [7, 44, 42], biodiversity indices [46], and algorithms that consider PD with ecological dependencies [14, 21, 27, 32, 37, 38]. In particular, since it is at the moment not clear which variant of phylogenetic diversity on networks is most useful in practice, it would be helpful to have them all in the same software tool so they can be compared and combined easily.

The implementation of Algorithm 1 demonstrates excellent scalability, enabling the maximization of phylogenetic diversity for large networks (see Figure 5). Our experiments also provide evidence for the value of scanwidth as a practical parameter for algorithm design in phylogenetics, showing that the parameter grows (much) slower than the level of a network (see Figure 4), a trend also noted in [22]. Combined with the relative elegance of Algorithm 1 compared to the theoretical contributions from [7, 24, 25], this suggests that scanwidth strikes a good balance between simplicity and tractability. Hence, it may be meaningful to explore whether a *semi-directed scanwidth* can be defined to help solve MapSPD in practice. Other interesting theoretical questions that could be useful to handle empirical data are whether the greedy approximation from [7] can be extended to the semi-directed setting, and whether ultrametricity can be exploited algorithmically.

From a practical point of view, our analysis of the *Xiphophorus* genus shows how `PaNDA` can offer a new perspective on biodiversity prioritization in the presence of reticulate events. It highlights a striking contrast with several traditional biodiversity indices, such as the *Shapley value* [18] and the *Fair Proportion Index* [23, 35], which basically measure the average contribution to phylogenetic diversity across all possible subsets of species. Such indices may rank two closely related species highly due to their individual contributions to diversity [46]. When prioritizing conservation of the top *k* highest ranked species [23], this can lead to suboptimal selections, since selecting two species that share a large portion of their ancestry likely yields diminishing returns. In contrast, using `PaNDA` to solve MapPD for a fixed value of *k* provides a different view that also takes this interdependence into account.

## Software and Data Availability

Supporting scripts, data sets and the source code of `PaNDA` are available at https://github.com/nholtgrefe/panda.

## Acknowledgment

We thank the anonymous reviewers of RECOMB 2026 for their detailed and constructive reviews. Their expertise and thoughtful suggestions substantially improved this work.

## A Appendix

### A.1 NP-hardness proof

**Theorem 1**. MapSPD *is* NP*-hard, even for binary semi-directed networks*.

*Proof*. We reduce from MapPD, which is NP-hard even for binary networks [7].

*Reduction*. Let ℐ = (𝒩^+^, *X, ω, k, D*) be an instance of MapPD with 𝒩^+^ being binary and having root *ρ*. Construct an instance ℐ^*′*^ = (𝒩 ^−^, *X*^*′*^, *ω*^*′*^, *k*^*′*^, *D*^*′*^) of MapSPD as follows. Let *M* be the sum of all edge weights plus one, and let *y* be a new taxon not in *X*. Set *X*^*′*^:= *X* ∪ {*y*}, *k*^*′*^:= *k* + 1 and *D*^*′*^:= *D* + *M*. Denote by 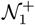 the directed network obtained from 𝒩^+^ by adding the leaf *y* and the edge (*ρ, y*). Set 𝒩 ^−^ to be the semi-directed network obtained from 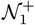, and set *ω*^*′*^(*e*):= *ω*(*e*) for all *e* ∈ *E*(𝒩^+^) with *ω*^*′*^((*ρ, y*)):= *M*.

*Correctness*. Since the out-degree of *ρ* is three, 𝒩 ^−^ can be obtained from 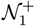 by only undirecting all non-reticulation edges. So, 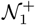 and 𝒩 ^−^ have the same vertex and edge sets (up to the undirecting of the non-reticulation edges). Thus, 𝒩 ^−^ is binary and *ω*^*′*^ is defined for all its edges. It remains to show the equivalence of ℐ and ℐ^*′*^. To this end, let *Y* ⊆ *X* be arbitrary. Then, we have that 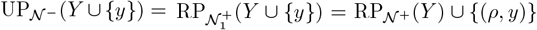. Because *ω*^*′*^((*ρ, y*)) = *M*, 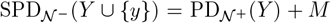. Since *M* being large forces *y* to be in any optimal solution for ℐ^*′*^, the equivalence follows from *D*^*′*^ = *D*+*M* and *k*^*′*^ = *k* + 1. Noting that the reduction takes polynomial time completes the proof.

### A.2 Proofs for Algorithm 1

#### Lemma 2

*Given an instance* (𝒩, *X, ω, k, D*) *of* MapPD *with* 𝒩 *nonbinary and a tree-extension* 𝒯_𝒩_ *of* 𝒩, *in* 𝒪(*m*) *time one can compute an equivalent instance* (𝒩 ^*′*^, *X, ω*^*′*^, *k, D*) *of* MapPD *and a tree-extension* 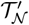 *of* 𝒩 ^*′*^ *such that* 𝒩 ^*′*^ *and* 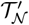 *are binary, m*^*′*^ ∈ 𝒪 (*m*), *and* 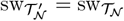.

*Proof*. A rooted binary tree *T* with leaf set {*x*_1_, …, *x*_*d*+1_} and *d* ≥ 1 is a *caterpillar tree* if the removal of all *x*_*i*_ results in a directed path *P* = (*v*_1_, …, *v*_*d*_), called the *spine*. For two leaves *x*_*i*_, *x*_*j*_ of *T* with in-neighbors *v*_*i*_, *v*_*j*_, respectively, we say that *x*_*i*_ *comes before x*_*j*_ if *i* ≤ *j*.

*Construction*. We let 𝒩 *′* be a carefully constructed *binary resolution* of 𝒩 and change 𝒯_𝒩_ into 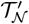 accordingly (see below). We set *ω*^*′*^(*e*):= *ω*(*e*) for all *e* ∈ *E*(𝒩) and *ω*^*′*^(*e*):= 0 for all *e* ∉ *E*(𝒩).

In particular, let *v*_1_ = *v* be a vertex with *d* + 1 ≥ 3 out-neighbors *w*_1_, …, *w*_*d*+1_ in 𝒩. Without loss of generality, assume that the *w*_*i*_ are ordered such that consecutive vertices belong to the same connected component of 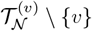, i.e., they are grouped according to the branch of 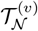 they are in. Let *T* be a caterpillar tree with root *v*_1_, leaf set {*w*_1_, …, *w*_*d*+1_}, spine *P* = (*v*_1_, …, *v*_*d*_) and the property that *w*_*i*_ comes before *w*_*j*_ in *T* if *i < j*. To obtain 𝒩 *′* from 𝒩, we replace *v*_1_ and its outgoing edges (*v*_1_, *w*_*i*_) in 𝒩 by *T*. To obtain 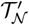 from 𝒯_𝒩_, we replace *v*_1_ by *P* such that the outgoing edges (*v*_1_, *z*_*j*_) of *v*_1_ in 𝒯_𝒩_ are replaced by edges (*v*_*d*_, *z*_*j*_). Similarly, let *v* = *v*_1_ be a vertex with *d* + 1 ≥ 3 in-neighbors *w*_1_, …, *w*_*d*+1_ in 𝒩. Let *T* ^*′*^ be a caterpillar tree with root *v*_1_, leaf set {*w*_1_, …, *w*_*d*+1_} and spine *P* ^*′*^ = (*v*_1_, …, *v*_*d*_). Let *T* be the graph obtained from *T* ^*′*^ by reversing all edge directions. To obtain 𝒩 *′* from 𝒩, we replace *v* and its incoming edges in 𝒩 by *T*. To obtain 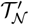 from 𝒯_𝒩_, we replace *v*_1_ by the directed path *P* = (*v*_*d*_, …, *v*_1_) such that the incoming edge (*z, v*_1_) of *v*_1_ in 𝒯_𝒩_ is replaced by (*z, v*_*d*_).

*Correctness*. Let *e* ∈ *E*(𝒩) be arbitrary. Then, note that our construction ensures that *e* is on a root-leaf path to a leaf *x* ∈ *X* in 𝒩 if and only if it is so in 𝒩 *′*. Since the new edges have a weight of zero, we obtain that PD_𝒩_*′* (*Y*) = PD_𝒩_ (*Y*) for all *Y* ⊆ *X*. Hence, the two instances are equivalent. Moreover, the construction can be done in a single traversal of the network and tree-extension, resulting in a running time of 𝒪 (*m*).

For the remaining conditions, first note that *m*^*′*^ is clearly ∈ 𝒪 (*m*). Since 𝒩 *′* is a valid binary resolution of 𝒩, it is trivially binary. By [6, Lem. 5], we may also assume that 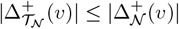 for all *v* ∈ *V* (𝒩), and hence our construction ensure that 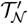 is binary. Next, observe that replacing vertices in 𝒯_𝒩_ that have a high in-degree in 𝒩 by a path preserves the property 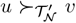 if *u* ≻_𝒩_*′ v*. Similarly, our construction accounting for the different branches of the 𝒯_𝒩_ ensures the same property is preserved when replacing vertices with high out-degree in 𝒩. Hence, 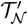 is a tree-extension of 𝒩*′*. Furthermore, we have that 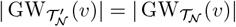 for all *v* ∈ *V* (𝒯_𝒩_). Each new vertex 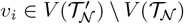 is part of a path *P*, starting at an existing vertex *v* ∈ *V* (𝒯_𝒩_). These vertices have the property 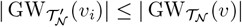 Hence, 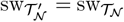.

#### Theorem 2

*Algorithm 1 solves instances* (𝒩, *X, ω, k, D*) *of* MapPD *in* 𝒪 (2^sw^ sw ·*k*^2^*m*) *time, if a tree-extension* 𝒯_𝒩_ *of* 𝒩 *of scanwidth* sw *is given*.

*Proof. Correctness*. Clearly, Algorithm 1 returns the correct result on Lines 22 to 24 if DP stores the intended values as described above. That this is true for the base cases on Lines 6, 8 and 10 of the algorithm can easily be seen. It remains to show the correctness of the recurrences on Lines 13, 15, 19 and 21. Let *v* ∈ *V* (𝒩) \ *X*, Φ ⊆ GW(*v*) and *t* ∈ [*k*]_0_ be arbitrary.

We use the shorthand notation 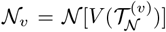 and *L*(*v*, Φ, *t*) = {*A* ⊆ *X* | *A* ⊆ *V* (𝒩_*v*_), off(*e*) ∩ *A* ≠ ∅ ∀*e* ∈ Φ, |*A*| = *t*}. The table stores the correct value if DP[*v*, Φ, *t*] stores the maximum value of 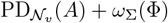 for any *A* ∈ *L*(*v*, Φ, *t*). We note that if there is no such set, then the value is −∞.

Since the recurrence on Line 19 is a simplification of the one on Line 21, we only prove correctness of the latter and assume that *v* has two children *u* and *w* in 𝒯_𝒩_. We show that, if Φ does not contain incoming edges, then *L*(*u*, Φ∩GW(*u*), *t*^*′*^) and *L*(*w*, Φ∩GW(*w*), *t*−*t*^*′*^) are a disjoint union of *L*(*v*, Φ, *t*), for some *t*^*′*^ ∈ [*t*]_0_. It immediately follows that Line 21 is correct. For each *A* ∈ *L*(*v*, Φ, *t*) we have *A*∩*V* (𝒩_*u*_) ∈ *L*(*u*, Φ ∩ GW(*u*), |*A* ∩ *V* (𝒩_*u*_)|). Since off(*e*) ∩ *A* ≠ ∅ for each *e* ∈ Φ, each *e* ∈ Φ ∩ GW(*u*) must have offspring in *A* ∩ *V* (𝒩_*u*_). Analogously, we get *A* ∩ *V* (𝒩_*w*_) ∈ *L*(*w*, Φ ∩ GW(*w*), |*A* ∩ *V* (𝒩_*w*_)|). As *u* and *w* are in children of *v* in 𝒯_𝒩_, the sets of vertices in the networks 𝒩_*u*_ and 𝒩_*w*_ are disjoint. This shows the first direction. Now, when *A*_*u*_ ∈ *L*(*u*, Φ ∩ GW(*u*), *t*^*′*^) and *A*_*w*_ ∈ *L*(*w*, Φ ∩ GW(*w*), *t* − *t*^*′*^) for some *t*^*′*^ ∈ [*t*]_0_, then *A*_*u*_ and *A*_*w*_ are disjoint and in total have *t* items. Further, by definition, off(*e*) ∩ (*A*_*u*_ ∪ *A*_*w*_) is non-empty for all *e* ∈ Φ. So, *A*_*u*_ ∪ *A*_*w*_ ∈ *L*(*v*, Φ, *t*).

Now, let *v* be the root, or let Φ contain an incoming edge of *v*. Since the recurrence on Line 13 is a simplification of the one on Line 15, we only prove correctness of the latter and assume that *v* has two children *u* and *w*. As induction hypothesis, suppose that DP stores the intended values for *u* and *w* in 𝒯_𝒩_. It is sufficient to show that DP[*v*, Φ, *t*] stores *d* ∈ ∈_≥0_ implies that there is an *A* ∈ *L*(*v*, Φ, *t*) with 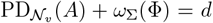 (that is, the table stores *at most* the maximum); and DP[*v*, Φ, *t*] stores at least 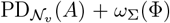 for each *A* ∈ *L*(*v*, Φ, *t*) (that is, the table stores *at least* the maximum). We handle the case that *L*(*v*, Φ, *t*) is empty—and therefore DP[*v*, Φ, *t*] stores −∞—independently.

First, suppose that DP[*v*, Φ, *t*] stores −∞. Then, for all *t*^*′*^ and all non-empty Ψ ⊆ Δ^−^(*v*), one of the tables DP[*u*, (Φ ∪ Ψ) ∩ GW(*u*), *t*^*′*^] or DP[*w*, (Φ ∪ Ψ) ∩ GW(*w*), *t* − *t*^*′*^] stores −∞. By the induction hypothesis, *L*(*u*, (Φ∪Ψ)∩GW(*u*), *t*^*′*^) or *L*(*w*, (Φ∪Ψ)∩GW(*w*), *t*−*t*^*′*^) is empty and therefore also *L*(*v*, Ψ, *t*). Now, let DP[*v*, Φ, *t*] store *d* ∈ ∈_≥0_. By the recurrence on Line 15 and the induction hypothesis, there is some Ψ ⊆ Δ^+^(*v*) and some *t*^*′*^ ∈ [*t*]_0_ such that there exist sets *A*_*u*_ ∈ *L*(*u, F*_*u*_, *t*^*′*^) and *A*_*w*_ ∈ *L*(*w, F*_*w*_, *t* − *t*^*′*^), where *F*_*u*_ = (Φ ∪ Ψ) ∩ GW(*u*) an *F*_*w*_ = (Φ ∪ Ψ) ∩ GW(*w*). Moreover,

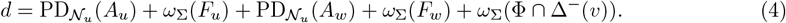

Let *A* = *A*_*u*_ ∪ *A*_*w*_. We show that *A* satisfies *A* ∈ *L*(*v*, Φ, *t*) and 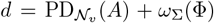. Since *A*_*u*_ ∈ *L*(*u, F*_*u*_, *t*^*′*^) and *A*_*w*_ ∈ *L*(*w, F*_*w*_, *t* − *t*^*′*^), we know *A* ⊆ *V* (𝒩_*v*_) = *V* (𝒩_*u*_) ∪ *V* (𝒩_*w*_). We conclude with *A*_*u*_ and *A*_*w*_ being disjoint that |*A*| = *t* and that 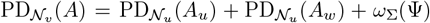. Because Φ = (Φ ∩ Δ^−^(*v*)) ∪ ((*F*_*v*_ ∪ *F*_*w*_) \ Ψ) and (Φ ∩ Δ^−^(*v*)), *F*_*v*_, and *F*_*w*_ are pairwise disjoint, we conclude 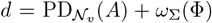. It remains to show that off(*e*) ∩ *A*≠ ∅ for each *e* ∈ Φ. Let *e* be an incoming edge of *v*. Then, since Ψ is non-empty, there is an edge *e*^*′*^ ∈ Ψ. Recall that *e*^*′*^ consequently is outgoing of *v* and off(*e*^*′*^) ⊆ off(*e*). Then, without loss of generality, let *e*^*′*^ be in *F*_*u*_. In consequence, off(*e*^*′*^) ∩ *A*_*u*_ is non-empty and therefore also off(*e*) ∩ *A*_*u*_ is non-empty. Now, let *e* ∈ Φ not be an incoming edge of *v*. Without loss of generality, *e* ∈ *F*_*u*_ and then off(*e*) ∩ *A*_*u*_ ∅, because *A*_*u*_ ∈ *L*(*u, F*_*u*_, *t*^*′*^).

Now, let there be an *A* ∈ *L*(*v*, Φ, *t*). If such a set does not exist then DP[*v*, Φ, *t*] stores −∞ by definition. Define *A*_*u*_ and *A*_*w*_ as the intersection of *A* with *X*(𝒩_*u*_):= *X* ∩ 𝒩_*u*_ and *X*(𝒩_*w*_):= *X* ∩ 𝒩_*w*_, respectively. These two sets are disjoint, as *X*(𝒩_*u*_) and *X*(𝒩_*w*_) are disjoint. Define Ψ to contain the outgoing edges *e*^*′*^ of *A*, such that off(*e*^*′*^) ∩ *A*≠ ∅. Because Φ has incoming edges of *v*, there is offspring of *v* in *A* and so Ψ is non-empty. We define, as before, *F*_*u*_:= (Ψ∪Φ) ∩GW(*u*) and *F*_*w*_ analogously. As we observe that *A*_*u*_ ∈ *L*(*u, F*_*u*_, |*A*_*u*_|) and *A*_*w*_ ∈ *L*(*w, F*_*w*_, |*A*_*w*_|), we conclude with the induction hypothesis that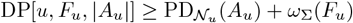 and 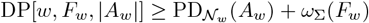. Then

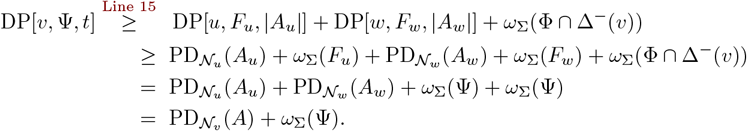

This completes the correctness proof.

*Running time*. By Lemma 2, we may assume that 𝒩 and 𝒯_𝒩_ are binary and instead show a running time of 𝒪 (2^sw^ sw ·*k*^2^*n*). Since | GW(*v*)| ≤ sw for all *v* ∈ *V* (𝒩), our table’s size is at most |*V* (𝒩)| · 2^sw^ · (*k* + 1). For each of these table entries, we perform a maximization over at most 3 · (*k* + 1) · 4 terms, since we assumed 𝒩 is binary. Thus, the time complexity becomes 𝒪 (2^sw^ sw ·*k*^2^ · *n*), as desired.

### A.3 Proofs for Algorithm 2

#### Lemma 3

*Rule 0 is correct and can be applied exhaustively in* 𝒪 (*m*) *time*.

*Proof*. Observe that an edge in a 2-blob contributes to the diversity of a set of taxa only if it is on an up-down path between two selected taxa. In that case, all of the edges in the blob, and the two incident cut-edges, are on such an up-down path. This proves correctness. The time complexity follows from the fact that finding and removing the 2-blobs can be done exhaustively in 𝒪 (*m*) time with a traversal of the network.

#### Lemma 4

*Rule 1 is correct and can be applied exhaustively in* 𝒪 (*nk*^2^) *time*.

*Proof*. Let ℐ = (𝒩, *X, f, ω, k, D*) denote an instance of G-MapSPD before the application of a reduction rule, and ℐ*′* = (𝒩 *′, X*^*′*^, *f* ^*′*^, *ω*^*′*^, *k, D*) the instance after the application.

It is easily visible that if the original instance ℐ has a solution *σ*, then *σ*^*′*^, where values of *X* \ {*x, y*} remain unchanged and *σ*^*′*^(*z*):= *σ*(*x*) + *σ*(*y*), is a solution for ℐ*′*. Conversely, from a solution *σ*^*′*^ of ℐ*′*, we can define a solution for ℐ by setting 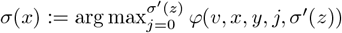 and *σ*(*y*):= *σ*^*′*^(*z*) − *σ*(*x*).

Every application of Rule 1 removes one leaf and takes 𝒪 (*k*^2^) time.

#### Lemma 5

*Rule 2 is correct and can be applied exhaustively in* 𝒪 (*n*) *time*.

*Proof*. Observe that only if leaves on both ends are selected, then one of the edges counts towards the diversity, and in that case, both. This shows the correctness. Further, this rule can be applied once per edge and each time in constant time. The number of edges that are on an edge-path to a leaf is 𝒪 (*n*).

#### Lemma 6

*A tree is reduced to a single leaf in* 𝒪 (*nk*^2^) *time, by exhaustively applying Rules 1 to 3*.

*Proof*. We prove that applying Rules 1 and 2 exhaustively is sufficient to correctly reduce trees to a single edge. Rule 3 then reduces the single edge to a single leaf in 𝒪 (1) time. Its correctness can be shown analogously to Lemma 4.

Let 𝒯 be a tree. We prove by induction on the number of leaves *n*, where *n* = 2 is trivial.

Let *n* ≥ 3 and the claim be correct for all trees with *n* − 1 leaves. Any tree on *n* leaves contains a cherry, that is, two leaves *x* and *y* that share a common neighbor *v*, or a degree-2 vertex *w*. If the latter is true, apply Rule 2 exhaustively. Afterward, we can apply Rule 1 to remove one of the leaves. The resulting graph is a tree on at most *n* − 1 vertices, and the claim follows by the induction hypothesis.

#### Lemma 7

*Let* ℐ = (𝒩, *X, f, ω, k, D*) *be an instance of* G-MapSPD *in which we can not apply any of the Rules 1 to 3. If* 𝒩 *contains a non-trivial blob B, then there exists a vertex v*_*B*_ ∈ *V* (*B*) *which is above every vertex of B. In particular, no reticulations can be above such vertices v*_*B*_.

*Proof*. Let *r* be a reticulation in *B* for which there are no reticulations above *r* (i.e., no reticulations have a semi-directed path to *r*). Let *u* be one of the in-neighbors of *r*. By Corollary 1 of [20], there is an up-down path between *u* and all vertices in *V* (*B*). By choice of *r*, there are no reticulations above *u*, so all such up-down paths must be a semi-directed path from *u*. This proves the existence part of the lemma.

To see the second part, let *v*_*B*_ ∈ *V* (*B*) be a vertex above every vertex of *B*. Suppose for a contradiction that there is a reticulation *r* above *v*_*B*_. Then there is a semi-directed path from *r* to *v*_*B*_. Since *v*_*B*_ is above *r*, there is a semi-directed path which is not an edge-path from *v*_*B*_ to *r*. Combining these two paths gives a semi-directed cycle, which is a cycle that can be oriented as a directed cycle. But this is not allowed by Theorem 2 of [20], which gives the required contradiction.

#### Lemma 8

*Let* ℐ = (𝒩, *X, f, ω, k, D*) *be an instance of* G-MapSPD *in which we can not apply any of the Rules 1 to 3. If* 𝒩 *contains a non-trivial blob, then a lowest blob of* 𝒩 *can be found in* 𝒪 (*m*) *time, where m is the number of edges in* 𝒩.

*Proof*. Suppose first that 𝒩 contains one blob *B*. Clearly, all neighbors of vertices of *B*, which are not in *V* (*B*), are leaves. By Lemma 7, there exists a vertex which is above every vertex of *B*. By definition, this is a designated vertex of *B*, and hence *B* is a lowest vertex.

Now suppose that 𝒩 contains at least two blobs. Define *v* ≥ *B* for a vertex *v* and a blob *B*, if there is a semi-directed path from *v* to *every* vertex of *B*. Define *B*_1_ ≥ *B*_2_ for blobs *B*_1_ and *B*_2_, if *v* ≥ *B*_2_ for some *v* ∈ *V* (*B*_1_), and if for no vertices *u V* ∈ (*B*_2_), *u ≥ B*_1_ holds.

Let *B* be a blob of 𝒩 which is smallest with respect to this partial ordering ≥, which has a neighboring blob *B*^+^ with *B*^+^ ≥ *B*. If there is no such blob, then let *B* be any blob of 𝒩. If *B* has at most one vertex with a non-leaf neighbor not in *B*, then we continue our argument in the next paragraph; otherwise, *B* must contain two vertices *v*_1_, *v*_2_ with non-leaf neighbors *u*_1_, *u*_2_ ∉ *V* (*B*) from blobs *B*_1_, *B*_2_, respectively. Observe that *v*_*i*_ must have a semi-directed path to all vertices in *V* (*B*_*i*_) for at least one of *i* = 1, 2, as otherwise, *V* (*B*) ∪ *V* (*B*_1_) ∪ *V* (*B*_2_) is part of a bigger blob. Without loss of generality, suppose *v*_1_ has a semi-directed path to all vertices in *V* (*B*_1_). We continue this process on *B*_1_; note that we can avoid returning to blobs that have already been visited, for example, by updating the partial ordering ≥ with *B* ≥ *B*_1_.

Since 𝒩 is a finite graph, such a process must terminate. We obtain a smallest blob *B*^∗^ in 𝒩 where at most one vertex *v* of *B*^∗^ has one non-leaf neighbor that is not in *V* (*B*)^∗^. We claim that *v* is above every vertex of *B*^∗^, thereby showing that *v* is a designated vertex of *B*^∗^, and consequently that *B*^∗^ is a lowest blob. Since *B*^∗^ is smallest, no reticulation of *B*^∗^ is above *v*. By Lemma 7, there exists a vertex *u* in *V* (*B*)^∗^ which is above every vertex in *V* (*B*)^∗^. Note that there must be an edge-path between *u* and *v*. Otherwise, there would be a reticulation in *V* (*B*)^∗^ above either *u* or *v*, which is not possible. But this means that *v* must also be above every vertex in *V* (*B*)^∗^, by appending the edge-path from *v* to *u* to the semi-directed paths from *u* to the other vertices, possibly removing duplicate edges. So *B*^∗^ must be a lowest blob with designated vertex *v*.

To show the runtime, note that one can traverse through the blobs as described above by visiting each edge at most once. Therefore, the lowest blob can be found in 𝒪 (*m*) time.

#### Lemma 9

*Let* 𝒩 *be a semi-directed network with a lowest blob B with designated vertex v*_*B*_ *and let P be a set of visible reticulations in V* (*B*). *Let w*, 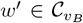 *be vertices with outgoing edges wr and w*^*′*^*r*^*′*^ *respectively, where r and r*^*′*^ *are reticulations with a directed path to an element in P*. *The set of vertices and edges on up-down paths between two vertices in P* ∪ {*w*} *and those in P* ∪ {*w*^*′*^} *are equivalent*.

*In other words, given a set of visible reticulations P*, *the definitions of V*_(*P*,1)_ *and E*_(*P*,1)_ *are independent of the vertex w of* 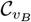 *which is chosen*.

*Proof*. If *P* = ∅ or if 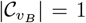, the claim follows immediately. So assume *P*≠ ∅ and 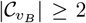. Let 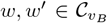 be two vertices with outgoing edges *wr* and *w*^*′*^*r*^*′*^, where *r* and *r*^*′*^ are reticulations with a semi-directed path to a reticulation *p* and *p*^*′*^ in *P*, respectively. We require *w* ≠ *w*^*′*^, but the others do not need to be distinct. We show that up-down paths of *U*_*w*_*′*:= UP(*P* ∪ {*w*^*′*^}) and *U*_*w*_:= UP(*P* ∪ {*w*}) cover the same set of vertices and edges.

First observe that any up-down path that has both endpoints in *P* is in both sets, *U*_*w*_*′* and *U*_*w*_.

Let *Q*_*w*+*w*_*′* be the edge-path from *w* to *w*^*′*^. Now let *Q* ∈ *U*_*w*_*′* use the edge *w*^*′*^*r*^*′*^, then *Q*_*w*+*w*_*′ Q* is a path in *U*_*w*_ that covers the set of vertices and edges of *Q*. Let *Q* ∈ *U*_*w*_*′* be an up-down path with *Q*_*w*+*w*_*′* ⊆ *Q*. We observe that *Q \ Q*_*w*+*w*_*′* is in *U*_*w*_ and that *Q*_*w*+*w*_*′ Q*_*w*_*′*_,*p*_*′* is in *U*_*w*_, where *Q*_*w*_*′*_,*p*_*′* is a directed path from *w*^*′*^ to *p*^*′*^ containing *w*^*′*^*r*^*′*^. These two paths cover the set of vertices and edges of *Q* For other updown paths *Q* ∈ *U*_*w*_*′*, add *Q*_*w*+*w*_*′* and remove doubled parts. Since *Q*_*w*+*w*_*′* is already covered, this shows that all vertices and edges that are covered with up-down paths in *U*_*w*_*′* are covered with up-down paths in *U*_*w*_.

#### Lemma 10

*The tree T*_(*P,α*)_ *can be constructed in* 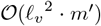 *time, where ℓ*_*v*_ *is the number of visible reticulations and m*^*′*^ *the number of edges in the original blob*.

*Proof*. For every pair of vertices in *P*_*α*_, we first compute the set of edges and vertices on up-down paths between them in 𝒪 (*m*^*′*^) time. There are 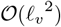 many pairs, so, finding *V*_(*P,α*)_ and *E*_(*P,α*)_ takes 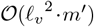 time. The remaining operations can all be done by a constant number of traversals of the blob, each taking 𝒪 (*m*^*′*^) time, thus proving the lemma.

#### Lemma 11

*Let* 𝒩 *be a semi-directed network with a lowest blob B with a designated vertex v*_*B*_. *Let x be a leaf neighbor of B. Exactly one of the following two statements is true*.

- *x has an edge-path to v*_*B*_.
- *x has an edge-path to a reticulation r. The reticulation r is visible with respect to x*.

*Proof*. By Condition (III) of Theorem 2 in [20], there can be no edge-path between two reticulations in a semi-directed network. Hence, there is at most one edge-path from *x* to a reticulation in 𝒩.

Suppose first that *x* has no such edge-path. We claim that *x* has an edge-path to *v*_*B*_. If not, then every semi-directed path from *v*_*B*_ to *x* contains a reticulation. Such a path must exist since *v*_*B*_ is above every vertex in *V* (*B*) by definition. But then *x* has an edge-path to a reticulation, which is a contradiction.

Now suppose that *x* has one edge-path to a reticulation in 𝒩, and call this reticulation *r*. Take any vertex *u* above *r*. We claim that every semi-directed path from *u* to *x* contains *r*. Suppose, for a contradiction, that this is not the case. Then we split into two cases.

Case 1: There exists a semi-directed path from *u* to *x* that is not an edge-path, which does not contain *r*. Find the last reticulation *r*^*′*^ on this path in the traversal order. The leaf *x* has an edge-path both to *r* and to *r*^*′*^, contradicting our assumption.

Case 2: There exists an edge-path from *u* to *x*. Since there is also an edge-path from *x* to *r*, we can combine a subset of the edges on these two edge-paths to obtain an edge-path from *u* to *r*. By definition, there is a semi-directed path from *u* to *r* which contains at least one reticulation edge. But this gives a semi-directed cycle, a cycle of directed edges and undirected edges which can be oriented to obtain a directed cycle, containing *u* and *r*. This is not possible by Condition (II) of Theorem 2 in [20], which gives our required contradiction.

So in particular, for any leaf *y* above *r*, all up-down paths between *x* and *y* must contain *r*. Thus *r* is visible with respect to *x*. As *v*_*B*_ is above *r*, there also cannot be an edge-path between *x* and *v*_*B*_.

#### Theorem 3

*Algorithm 2 solves instances* (𝒩, *X, f, ω, k, D*) *of* G-MapSPD *in* 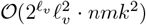 *time, where ℓ*_*v*_ *is the visible vertex level, n is the number of vertices, and m is the number of edges in* 𝒩.

*Proof*. By Lemma 3, we can preprocess the network such that there are no 2-blobs, and that there are thus 𝒪 (*n*) blobs in total.

We now prove the correctness of Algorithm 2 with three claims. Let ℐ = (𝒩, *X, f, ω, k, D*) be an instance of G-MapSPD.

#### Claim 12

*If* 𝒩 *is a tree, then Algorithm 2 correctly returns whether* ℐ *is a* `yes`*-instance of* G-MapSPD *in Line 4 or Line 5*.

After Line 1, all subtrees in 𝒩 are reduced to leaves. Thus, if 𝒩 is a tree, we return `yes` or `no` in Line 4 or 5. The correctness of Claim 12 follows with Lemma 6.

Now let 𝒩 be a network which has at least one reticulation and on which we can not apply Rules 1 to 3. Let *B* denote the lowest blob found in Line 6—and to which the algorithm is applied to until a return—with designated vertex *v*_*B*_, and let *X*_*B*_ denote the leaf neighbors of *B*.

#### Claim 13

*There is a solution σ for* ℐ *which only spends budget on (a subset of) X*_*B*_ *if and only if Algorithm 2 returns* `yes` *in Line 11*.

Finally, let Algorithm 2 not return in Line 4, 5, or 11. Let ℐ*′* = (𝒩^*′*^, *X*^*′*^, *f* ^*′*^, *ω*^*′*^, *k, D*) be the instance given into the recursion in Line 21.

#### Claim 14

ℐ *is a* `yes`*-instance of* G-MapSPD *if and only if* ℐ*′ is a* `yes`*-instance of* G-MapSPD.

We observe that these three claims ar sufficient to prove the correctness of Algorithm 2. We continue proving Claim 13 and Claim 14, afterward.

*Proof of Claim 13*. Let *σ* be a solution for ℐ which only spends a budget of *k* on *X*_*σ*_ ⊆ *X*_*B*_. Let *α* be 1 if *v* has an edge-path to a leaf in *X*_*σ*_, and let *α* be 0, otherwise. Let *P* be the set of reticulations which have an edge-path to a vertex in *X*_*σ*_. Since each taxon has an edge-path to at most one reticulation (Lemma 11), the set *P* is well-defined. We show that in tree *T*:= *T*_(*P,α*)_, the diversity of *X*_*σ*_ is PD_𝒩_ (*X*_*σ*_) − *H*_(*P,α*)_ − *ω*_Σ_(*E*_(*P,α*)_) + *m*_(*P,α*)_ · *M*. We implicitly use that *P*_1_ is well-defined (Lemma 9). In the light of Lemma 6, this is sufficient to prove that in Line 10 a `yes` is triggered. We consider the steps of creating *T*. In Step 1, we remove vertices in hidden(*V*_(*P,α*)_), which is equivalent to setting *σ* to 0 in every leaf in hidden(*V*_(*P,α*)_). We thus reduce the diversity by *H*_(*P,α*)_. Recall that *E*_(*P,α*)_ is the set of edges on up-down paths between vertices in *P*. As we can extend these paths on both ends to leaves in *X*_*σ*_, the weight of *E*_(*P,α*)_ is considered in PD_𝒩_ (*X*_*σ*_), but compressed in Step 2. In Step 4, we add *m*_(*P,α*)_ edges of length *M* as connection between 𝒞_*r*_ and *v*_*P,α*_, which thus are all on paths between specific leaves in *X*_*σ*_. In Step 1 and 5, vertices are removed that are not on up-down paths between vertices in *X*_*σ*_. In consequence, `yes` is returned in Line 11. This completes this direction.

Conversely, let `yes` be returned in Line 11. Thus, *f* (*u*_(*P,α*)_, *k*) + *H*_(*P,α*)_ + *ω*_Σ_(*E*_(*P,α*)_) − *m*_(*P,α*)_ · *M* ≥ *D*, for some *P* ⊆ *R*_*V*_ and some *α* ∈ {0, 1}. With Lemma 6, we conclude that there is a function *σ* that spends a budget of *k* on leaves of *T*_(*P,α*)_ such that 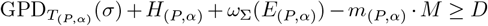. With similar arguments as before, we conclude that *σ* is a solution for 𝒩. ⋄

*Proof of Claim 14*. We can show this claim is relatively analogous to Claim 13 and thus only point out differences.

In Step 3 of creating *T*_(*P,α*)_, we add two leaf neighbors of *v* to ensure that there is always an up-down path to *v*, since we want to spend a positive budget on taxa in *X \ X*_*B*_. These are two leaves to create the possibility of spending a budget of *h ∈* {0, 1} on *X*_*B*_. Consequently, we have to allow for a budget of *h* + 2 in *u*_(*P,α*)_.

Instead of returning instantly, we search for each *h* the *P*, such that the diversity contribution in *B* is maximized. Which *P* ⊆ *R*_*V*_ was chosen best for a specific *h* is irrelevant in the end. ⋄

To show the running time, we recall that finding a lowest blob can be done in 𝒪 (*m*) time, Lemma 8, and Rule 0 can be applied exhaustively in 𝒪 (*m*) time, Lemma 3. Note that Rule 0 will only need to be called once before any loop or recursion, since the remainder of the algorithm creates no new 2-blobs.

We observe that the for loop in Line 8 is called 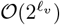 times, because |*R*_*v*_| ∈ 𝒪 (*ℓ*_*v*_). The trees *T*_(*P,α*)_ are computed in 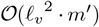 time each, by Lemma 10, where *m*^*′*^ is the number of edges in the blob used to create *T*_(*P,α*)_. Applying Rules 1 to 3 is done in 𝒪 (*nk*^2^) time, Lemma 6.

Each time Algorithm 2 is called, at least one blob is reduced, such that 𝒪 (*n*) are necessary. Thus, the overall running time is 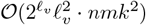.

## Footnotes

1 Recall that this is the subgraph of 𝒩 induced by 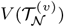.

2 Budgeted Max-PD [34] is a variant of Max-PD on unrooted phylogenetic trees in which each taxon *x* has a cost *c*_*x*_ of survival. It is a special case of G-MapSPD, if *C*—the biggest cost of a taxon—is polynomial in the input size. That is, by setting *f* (*x, h*) to −∞ for *h < c*_*x*_, and 0 otherwise. Budgeted Max-PD can be solved in 𝒪 (*k*^2^*n*) time [34], or in 𝒪 (*C*^2^*n*^3^) time [28]. However, Budgeted Max-PD is NP-hard, when taxa can have arbitrarily big costs. In that case, defining *f* is not possible in polynomial time.

3 In practice, choosing *k* = *n* yields a trivial solution. Nevertheless, it serves to illustrate the computability of our approach even for the largest possible values of *k*.

## References

[1] Elizabeth S. Allman, Hector Baños, John A. Rhodes, and Kristina Wicke. NANUQ+: A divide- and-conquer approach to network estimation. Algorithms for Molecular Biology, 20(1):14, 2025. doi:10.1186/s13015-025-00274-w.

[2] Hector Baños. Identifying Species Network Features from Gene Tree Quartets Under the Coalescent Model. Bulletin of Mathematical Biology, 81(2):494–534, 2019. doi:10.1007/s11538-018-0485-4.

[3] Eric Bapteste, Leo van Iersel, Axel Janke, Scot Kelchner, Steven Kelk, James O. McInerney, David A. Morrison, Luay Nakhleh, Mike Steel, Leen Stougie, and James Whitfield. Networks: expanding evolutionary thinking. Trends in Genetics, 29(8):439–441, 2013. doi:10.1016/j.tig.2013.05.007.

[4] Shalome A. Bassett, Wayne Young, Matthew P.G. Barnett, Adrian L. Cookson, Warren C. McNabb, and Nicole C. Roy. Changes in Composition of Caecal Microbiota Associated with Increased Colon Inflammation in Interleukin-10 Gene-Deficient Mice Inoculated with Enterococcus Species. Nutrients, 7(3):1798–1816, 2015. doi:10.3390/nu7031798.

[5] Paul Bastide, Claudia Solís-Lemus, Ricardo Kriebel, K. William Sparks, and Cécile Ané. Phylogenetic Comparative Methods on Phylogenetic Networks with Reticulations. Systematic Biology, 67(5):800–820, 2018. doi:10.1093/sysbio/syy033.

[6] Vincent Berry, Celine Scornavacca, and Mathias Weller. Scanning Phylogenetic Networks Is NPhard. In Proceedings of the 46th International Conference on Current Trends in Theory and Practice of Computer Science (SOFSEM 2020), pages 519–530. Springer, 2020. doi:10.1007/978-3-030-38919-2\_42.

[7] Magnus Bordewich, Charles Semple, and Kristina Wicke. On the complexity of optimising variants of phylogenetic diversity on phylogenetic networks. Theoretical Computer Science, 917:66–80, 2022. doi:10.1016/j.tcs.2022.03.012.

[8] Eduardo Sonnewend Brondízio, Josef Settele, Sandra Diaz, and Hien Thu Ngo. Global assessment report on biodiversity and ecosystem services of the Intergovernmental Science-Policy Platform on Biodiversity and Ecosystem Services. IPBES, 2019. doi:10.5281/zenodo.3831673.

[9] Cedoljub Bundalovic-Torma, Darrell Desveaux, and David S. Guttman. RecPD: A Recombinationaware measure of phylogenetic diversity. PLoS Computational Biology, 18(2):e1009899, 2022. doi:10.1371/journal.pcbi.1009899.

[10] Gabriel Cardona, Francesc Rosselló, and Gabriel Valiente. Extended Newick: it is time for a standard representation of phylogenetic networks. BMC Bioinformatics, 9(1):532, 2008. doi:10.1186/1471-2105-9-532.

[11] Rongfeng Cui, Molly Schumer, Karla Kruesi, Ronald Walter, Peter Andolfatto, and Gil G. Rosenthal. Phylogenomics reveals extensive reticulate evolution in Xiphophorus fishes. Evolution, 67(8):2166–2179, 2013. doi:10.1111/evo.12099.

[12] Reinhard Diestel. Graph Theory, volume 173. Springer Nature, 2025. doi:10.1007/978-3-662-70107-2.

[13] Daniel P. Faith. Conservation evaluation and phylogenetic diversity. Biological Conservation, 61(1):1–10, 1992. doi:10.1016/0006-3207(92)91201-3.

[14] Beáta Faller, Charles Semple, and Dominic Welsh. Optimizing Phylogenetic Diversity with Ecological Constraints. Annals of Combinatorics, 15(2):255–266, 2011. doi:10.1007/s00026-011-0093-6.

[15] Elizabeth Gross and Colby Long. Distinguishing Phylogenetic Networks. SIAM Journal on Applied Algebra and Geometry, 2(1):72–93, 2018. doi:10.1137/17m1134238.

[16] Rikki Gumbs, Claudia L. Gray, Monika Böhm, Ian J. Burfield, Olivia R. Couchman, Daniel P. Faith, Félix Forest, Michael Hoffmann, Nick J.B. Isaac, Walter Jetz, et al. The EDGE2 protocol: Advancing the prioritisation of Evolutionarily Distinct and Globally Endangered species for practical conservation action. PLoS Biology, 21(2):e3001991, 2023. doi:10.1371/journal.pbio.3001991.

[17] Rikki Gumbs, Claudia L. Gray, Monika Böhm, Michael Hoffmann, Richard Grenyer, Walter Jetz, Shai Meiri, Uri Roll, Nisha R. Owen, and James Rosindell. Global priorities for conservation of reptilian phylogenetic diversity in the face of human impacts. Nature Communications, 11(1):2616, 2020. doi:10.1038/s41467-020-16410-6.

[18] Claus-Jochen Haake, Akemi Kashiwada, and Francis Edward Su. The Shapley value of phylogenetic trees. Journal of Mathematical Biology, 56(4):479–497, 2008. doi:10.1007/s00285-007-0126-2.

[19] Niels Holtgrefe, Katharina T. Huber, Leo van Iersel, Mark Jones, Samuel Martin, and Vincent Moulton. Squirrel: Reconstructing Semi-directed Phylogenetic Level-1 Networks from Four-Leaved Networks or Sequence Alignments. Molecular Biology and Evolution, 42(4):msaf067, 2025. doi:10.1093/molbev/msaf067.

[20] Niels Holtgrefe, Katharina T Huber, Leo van Iersel, Mark Jones, and Vincent Moulton. Characterizing semi-directed phylogenetic networks and their multi-rootable variants. Theory in Biosciences, 145(1):4, 2026. doi:10.1007/s12064-025-00453-8.

[21] Niels Holtgrefe, Jannik Schestag, and Norbert Zeh. Limits of Kernelization and Parametrization for Phylogenetic Diversity with Dependencies. In Proceedings of the 17th Latin American Theoretical Informatics Symposium (LATIN 2026). Springer, 2026. 2602.12959.

[22] Niels Holtgrefe, Leo van Iersel, and Mark Jones. Exact and Heuristic Computation of the Scanwidth of Directed Acyclic Graphs. arXiv preprint, 2024. 2403.12734.

[23] Nick J.B. Isaac, Samuel T. Turvey, Ben Collen, Carly Waterman, and Jonathan E.M. Baillie. Mammals on the EDGE: Conservation Priorities Based on Threat and Phylogeny. PloS One, 2(3):e296, 2007. doi:10.1371/journal.pone.0000296.

[24] Mark Jones and Jannik Schestag. How Can We Maximize Phylogenetic Diversity? Parameterized Approaches for Networks. In Proceedings of the 18th International Symposium on Parameterized and Exact Computation (IPEC 2023), pages 30:1–30:12. Schloss-Dagstuhl-Leibniz Zentrum für Informatik, 2023. doi:10.4230/LIPIcs.IPEC.2023.30.

[25] Mark Jones and Jannik Schestag. Parameterized Algorithms for Diversity of Networks with Ecological Dependencies. In Proceedings of the 20th International Symposium on Parameterized and Exact Computation (IPEC 2025), pages 11:1–11:21. Schloss-Dagstuhl-Leibniz Zentrum für Informatik, 2025. doi:10.4230/LIPIcs.IPEC.2025.11.

[26] Joshua A. Justison, Claudia Solis-Lemus, and Tracy A. Heath. SiPhyNetwork: An R package for simulating phylogenetic networks. Methods in Ecology and Evolution, 14(7):1687–1698, 2023. doi:10.1111/2041-210x.14116.

[27] Christian Komusiewicz and Jannik Schestag. Maximizing Phylogenetic Diversity under Ecological Constraints: A Parameterized Complexity Study. In Proceedings of the 44th IARCS Annual Conference on Foundations of Software Technology and Theoretical Computer Science (FSTTCS 2024), pages 28:1–28:18. Schloss-Dagstuhl-Leibniz Zentrum für Informatik, 2024. doi:10.4230/LIPIcs.FSTTCS.2024.28.

[28] Christian Komusiewicz and Jannik Schestag. A Multivariate Complexity Analysis of the Generalized Noah’s Ark Problem. Discrete Applied Mathematics, 382:137–154, 2025. doi:10.1016/j.dam.2025.11.037.

[29] Sungsik Kong, Claudia Solís-Lemus, and George P. Tiley. Phylogenetic networks empower biodiversity research. Proceedings of the National Academy of Sciences, 122(31):e2410934122, 2025. doi:10.1073/pnas.2410934122.

[30] Samuel Martin, Niels Holtgrefe, Vincent Moulton, and Richard M. Leggett. Algebraic invariants for inferring 4-leaf semi-directed phylogenetic networks. Systematic Biology, page syaf071, 2025. doi:10.1093/sysbio/syaf071.

[31] Rafael Molina-Venegas, Miguel A. Rodriguez, Manuel Pardo-de Santayana, Cristina Ronquillo, and David J. Mabberley. Maximum levels of global phylogenetic diversity efficiently capture plant services for humankind. Nature Ecology & Evolution, 5(5):583–588, 2021. doi:10.1038/s41559-021-01414-2.

[32] Vincent Moulton, Charles Semple, and Mike Steel. Optimizing phylogenetic diversity under constraints. Journal of Theoretical Biology, 246(1):186–194, 2007. doi:10.1016/j.jtbi.2006.12.021.

[33] Fabio Pardi and Nick Goldman. Species Choice for Comparative Genomics: Being Greedy Works. PLoS Genetics, 1, 2005. doi:10.1371/journal.pgen.0010071.

[34] Fabio Pardi and Nick Goldman. Resource-Aware Taxon Selection for Maximizing Phylogenetic Diversity. Systematic Biology, 56(3):431–444, 2007. doi:10.1080/10635150701411279.

[35] David W. Redding. Incorporating Genetic Distinctness and Reserve Occupancy into a Conservation Prioritisation Approach. Master’s thesis, University of East Anglia, 2003.

[36] Dan F. Rosauer and Walter Jetz. Phylogenetic endemism in terrestrial mammals. Global Ecology and Biogeography, 24(2):168–179, 2015. doi:10.1111/geb.12237.

[37] Jannik Schestag. Weighted Food Webs Make Computing Phylogenetic Diversity So Much Harder. In Proceedings of the 51st International Conference on Current Trends in Theory and Practice of Computer Science (SOFSEM 2026), page 187–202. Springer, 2026.

[38] Jannik Schestag and Norbert Zeh. The First Known Problem That Is FPT with Respect to Node Scanwidth but Not Treewidth. 2026. 2602.06903.

[39] Claudia Solís-Lemus and Cécile Ané. Inferring Phylogenetic Networks with Maximum Pseudolikelihood under Incomplete Lineage Sorting. PLoS Genetics, 12(3):e1005896, 2016. doi:10.1371/journal.pgen.1005896.

[40] Claudia Solís-Lemus, Paul Bastide, and Cécile Ané. PhyloNetworks: A Package for Phylogenetic Networks. Molecular Biology and Evolution, 34(12):3292–3298, 2017. doi:10.1093/molbev/msx235.

[41] Mike Steel. Phylogenetic Diversity and the Greedy Algorithm. Systematic Biology, 54(4):527–529, 2005. doi:10.1080/10635150590947023.

[42] Leo van Iersel, Mark Jones, Jannik Schestag, Celine Scornavacca, and Mathias Weller. Average-Tree Phylogenetic Diversity of Networks. In Proceedings of the 25th International Workshop on Algorithms in Bioinformatics (WABI 2025), pages 14:1–14:21. Schloss Dagstuhl–Leibniz-Zentrum für Informatik, 2025. doi:10.4230/LIPIcs.WABI.2025.15.

[43] Leo van Iersel, Mark Jones, Jannik Schestag, Celine Scornavacca, and Mathias Weller. Average-Tree Phylogenetic Diversity Parameterized by Scanwidth and Invisibility. Manuscript Under Review, 2025.

[44] Leo van Iersel, Mark Jones, Jannik Schestag, Celine Scornavacca, and Mathias Weller. Phylogenetic Network Diversity Parameterized by Reticulation Number and Beyond. In Proceedings of the 22nd RECOMB International Workshop on Comparative Genomics (RECOMB-CG 2025), pages 107–130. Springer, 2025. doi:10.1007/978-3-031-94928-9_7.

[45] Logan Volkmann, Iain Martyn, Vincent Moulton, Andreas Spillner, and Arne O. Mooers. Prioritizing Populations for Conservation Using Phylogenetic Networks. PloS One, 9(2):e88945, 2014. doi:10.1371/journal.pone.0088945.

[46] Kristina Wicke and Mareike Fischer. Phylogenetic diversity and biodiversity indices on phylogenetic networks. Mathematical Biosciences, 298:80–90, 2018. doi:10.1016/j.mbs.2018.02.005.

